# Extensive admixture among karst-obligate salamanders reveals evidence of recent divergence and gene exchange through aquifers

**DOI:** 10.1101/2024.07.31.606001

**Authors:** Chris C. Nice, Katherine L. Bell, Zachariah Gompert, Lauren K. Lucas, James R. Ott, Ruben U. Tovar, Paul Crump, Peter H. Diaz

**Author notes:** Corresponding author: Chris Nice, Department of Biology, 601 University Drive, Texas State University, San Marcos, TX 78666, USA.

## Abstract

Karst ecosystems often contain extraordinary biodiversity, but the complex underground aquifers of karst regions present challenges for assessing and conserving stygobiont diversity and investigating their evolutionary history. We examined the karst-obligate salamanders of the *Eurycea neotenes* species complex in the Edwards Plateau region of central Texas using population genomics data to address questions about population connectivity and the potential for gene exchange within the underlying aquifer system. The *Eurycea neotenes* species complex has historically been divided into three nominal species, but their status, and spatial extent of species ranges, have remained uncertain. We discovered evidence of extensive admixture within the species complex and with adjacent lineages. We observed relatively low levels of differentiation among all sampling localities which supports the hypothesis of recent divergence. Nominal taxonomy, aquifer region and geography accounted for a modest amount of the overall population genomic variation, but these predictors were largely collinear and difficult to disentangle. Importantly, the taxonomy of the three nominal species does not reflect the admixture apparent in clustering analyses. Inference of migration events revealed a complex pattern of gene exchange, suggesting that *Eurycea* salamanders have a dynamic history of dispersal through the aquifer system. These results highlight the need for greater understanding of how stygobiont populations are connected via dispersal and gene exchange through karst aquifers.

## Introduction

Limestone karst landforms are considered “islands” of terrestrial and aquatic habitats that often exhibit extraordinary biodiversity (Barr Jr & Holsinger, 1985; Clements *et al*., 2006; Hutchins *et al*., 2021; Longley, 1981). Within karst islands, cave and aquifer systems form a complex, and often human-inaccessible, subterranean landscape for both terrestrial and aquatic organisms. The subterranean complexity contributes to species richness through dispersal limitation, multiple colonizations, opportunities for vicariance, and diverse, sometimes extreme habitats that foster the evolution of ecological, behavioral and morphological specialization (Bendik *et al*., 2013; Culver *et al*., 2009).

Karst biotas include many endemic species with relatively narrow geographic ranges that are also sensitive to perturbations. Conservation of these ecosystems and their constituent species is sometimes hampered by a lack of knowledge about their biogeographic, evolutionary and demographic histories. In particular: 1) The number of distinct lineages or species might be underestimated due to under sampling, insufficient taxonomic knowledge (e.g. Hutchins *et al*., 2021), or convergent or parallel evolution (Wiens *et al*., 2003); or diversity can be overestimated due to conflicts between morphological and genetic data. 2) The extent of species ranges (or the boundaries between them) are often quite difficult to delineate given subterranean complexity and inaccessibility in karst regions. As a result of the complexity of subterranean habitat geography, 3) the conduits for population connectivity (via gene flow) can be difficult to predict and might differ among even closely related lineages with divergent adaptive histories (Katz *et al*., 2018). Furthermore, the conduits for gene flow can be dynamic depending on aquifer recharge rates and water levels (Longley, 1981). We confront these knowledge gaps with population genomics data from extensive sampling of the *Eurycea neotenes* salamander species complex in the central Edwards Plateau region of Texas, one of the most diverse karst systems (Longley, 1981).

The Edwards-Trinity Aquifer system underlies the Edwards Plateau region of central Texas (Figure 1), an area of uplifted limestone that is home to a large number of cave- and spring-associated species. The rich stygobiont diversity of the aquifers is increasingly impacted by encroaching human population density in the area. Human impacts include altered recharge patterns (Lamichhane & Shakya, 2019), higher levels of nutrients (Boyer *et al*., 2002) and contaminants (Diaz *et al*., 2020), as well as lowered water tables in parts of the aquifer (Green *et al*., 2011). These anthropogenic stressors are exacerbated by climate change and have the potential to profoundly affect regional and local biodiversity.

**Figure 1:**
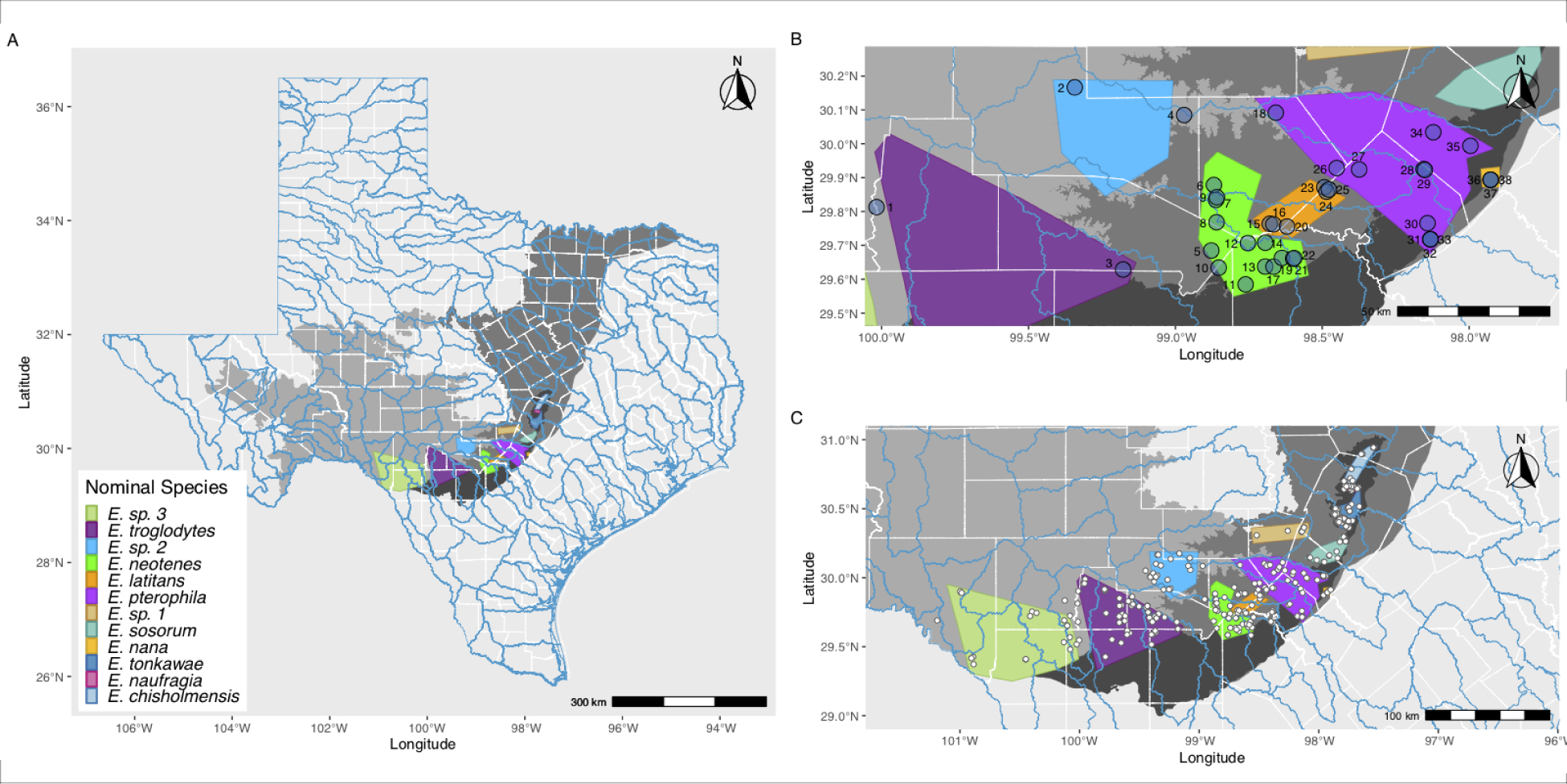
Map of the central Edwards Plateau region of Texas with *Eurycea* salamander locations, approximate ranges and sampling localities for this study indicated. A. Major aquifers in the state of Texas and approximate ranges of nominal *Eurycea* lineages. Shading indicates specific components of the aquifer system: Edwards-Trinity Aquifer (light gray), Trinity Aquifer (medium gray), Edwards Aquifer (dark gray). These aquifer components are variously connected and form the Edwards-Trinity aquifer system. The approximate ranges of the nominal *Eurycea* lineages are indicated by colored polygons. B. Focused map of sampling localities for this study. Locality numbers follow Table 1. C. Approximate locations of *Eurycea* occurrences, including known (historic) localities, localities sampled by Devitt et al. (2019) and sampling localities for this study. Range information based on Devitt et al. (2019), Chippindale et al. (2000), Bendik et al. (2013) and personal observations.

**Table 1:** Sampling details. Locality names and samples sizes (n) for individuals genotyped for each locality.

| Locality Number | Locality Name | n | Nominal Taxonomy | Major Aquifer |
| --- | --- | --- | --- | --- |
| 1 | Nueces 335 | 2 | *E. troglodytes* | Edwards-Trinity |
| 2 | Fessenden Springs | 23 | *E. sp. 2* | Edwards-Trinity |
| 3 | Hill Country SNA | 11 | *E. troglodytes* | Trinity |
| 4 | Western Kerr | 14 | *E. sp. 2* | Edwards-Trinity |
| 5 | Osborn Spring | 13 | *E. neotenes* | Trinity |
| 6 | Possum Creek Spring | 14 | *E. neotenes* | Trinity |
| 7 | Brown Ranch | 42 | *E. neotenes* | Trinity |
| 8 | Albert and Bessie Kronkosky SNA | 44 | *E. neotenes* | Trinity |
| 9 | Salamander Spring | 16 | *E. neotenes* | Trinity |
| 10 | Pecan Springs Cave | 1 | *E. neotenes* | Trinity |
| 11 | Government Canyon | 22 | *E. neotenes* | Edwards |
| 12 | Mueller's Ranch | 2 | *E. neotenes* | Trinity |
| 13 | Helotes Creek Spring | 3 | *E. neotenes* | Trinity |
| 14 | Maverick Ranch | 28 | *E. neotenes* | Trinity |
| 15 | Cascade Caverns | 28 | *E. latitans* | Trinity |
| 16 | Pfeiffer Ranch | 4 | *E. latitans* | Trinity |
| 17 | Leahs Spring | 9 | *E. neotenes* | Trinity |
| 18 | Peavey's Spring | 1 | *E. latitans* | Edwards-Trinity |
| 19 | Leon Springs | 29 | *E. neotenes* | Trinity |
| 20 | Badweather Pit | 3 | *E. latitans* | Trinity |
| 21 | Camp Bullis Stealth | 8 | *E. neotenes* | Trinity |
| 22 | Camp Bullis Sharon | 12 | *E. neotenes* | Trinity |
| 23 | Guadalupe River State Park | 22 | *E. latitans* | Trinity |
| 24 | Honey Creek SNA Springs | 32 | *E. latitans* | Trinity |
| 25 | Preserve Cave | 10 | *E. latitans* | Trinity |
| 26 | Sattler's Deep Pit | 2 | *E. latitans* | Trinity |
| 27 | Rebecca Spring | 2 | *E. latitans* | Trinity |
| 28 | Devil's Backbone | 18 | *E. pterophila* | Edwards |
| 29 | Ott's spring | 17 | *E. pterophila* | Trinity |
| 30 | Hueco Springs | 15 | *E. pterophila* | Edwards |
| 31 | Comal Spr Run One | 14 | *E. pterophila* | Edwards |
| 32 | Comal Spr Run Three | 19 | *E. pterophila* | Edwards |
| 33 | Comal SPr Spring Island | 22 | *E. pterophila* | Edwards |
| 34 | Jacobs Well | 21 | *E. pterophila* | Trinity |
| 35 | Fern Bank | 35 | *E. pterophila* | Trinity |
| 36 | San Marcos below dam | 18 | *E. nana* | Edwards |
| 37 | San Marcos Diversion | 16 | *E. nana* | Edwards |
| 38 | San Marcos Hotel | 15 | *E. nana* | Edwards |
| Total |  | 607 |  |  |

Among the Edwards Plateau region stygobionts impacted by these stressors are up to 14 species of endemic, fully aquatic, neotenic salamanders of the genus *Eurycea* (Plethodontidae: Hemidactyliinae) (Devitt *et al*., 2019) that inhabit springs and caves throughout the Edwards Plateau (Figure 1). Ten of these species are listed as threatened or endangered at the federal or state level (https://ecos.fws.gov, https://tpwd.texas.gov/). These salamanders are adapted to the aquifer habitats since their colonization of the aquifers and are quite specialized. Some of the species are adapted to deep aquifer habitats and exhibit classic convergent troglobitic (subsurface) morphologies that include loss of eyes, depigmentation, and elongation of limbs and skull, while other species reside in springs and caves and exhibit surface-dwelling morphologies, retaining eyes and pigmentation (Devitt *et al*., 2019; Hillis *et al*., 2001). The *Eurycea* salamanders of the Edwards Plateau are specialized stygobionts that are relatively intolerant of fluctuations in water temperature (Barr & Babbitt, 2002; Crow *et al*., 2016) and quality (Bowles *et al*., 2006). These salamanders are completely reliant upon groundwater, which makes the encroachment of urban areas and climate change increasing conservation concerns (Bendik *et al*., 2014; Diaz *et al*., 2020).

Despite considerable research into the phylogeny and population genetic structure of spring salamanders in central Texas (e.g. Bendik *et al*., 2013; Chippindale *et al*., 2000; Devitt *et al*., 2019; Hillis *et al*., 2001; Lucas *et al*., 2009), uncertainty remains regarding the boundaries between, and the extent of gene exchange among, some of the lineages. We focused on a segment of the diversity of *Eurycea* in the southeastern Edwards Plateau that is home to several very closely related nominal species in the *E. neotenes* species complex (Chippindale *et al*., 2000; Bendik *et al*., 2013; Devitt *et al*., 2019). Historically, four species have been recognized within the region. These include the Texas salamander (*E. neotenes*, Bishop & Wright, 1937), the Cascade Cavern’s salamander (*E. latitans*, Smith & Potter, 1946), the Fern Bank Salamander (*E. pterophila*, Burger *et al*., 1950) and the Comal Blind Salamander (*E. tridentifera*, Mitchell & Reddell, 1965), however, Devitt *et al*. (2019) synonomized *E. tridentifera* within *E. latitans*. This group has a complicated taxonomic history with morphological (Sweet, 1984) and genetic (Bendik *et al*., 2013; Devitt *et al*., 2019) evidence of incomplete isolation and ongoing gene flow. The uncertainty about these species and the extent of gene flow among them is due in part to the apparent recent evolution of the group and relatively limited sampling in the ranges of these salamanders (Chippindale *et al*., 2000; Devitt *et al*., 2019).

Here we sampled the *Eurycea neotenes* complex of salamanders from the Guadalupe and San Antonio river drainages in central Texas. We sampled extensively with respect to both geography and samples per locality and generated population genomics data to address the following questions: 1) do patterns of population genomic variation support the nominal taxonomy of three distinct lineages of surface salamanders corresponding to *E. pterophila*, *E. neotenes* and *E. latitans*?, 2) how differentiated are populations and lineages of these salamanders?, and 3) to what extent have populations or lineages of these salamanders been connected by historical or contemporary gene flow? The answers to these questions will provide critical insights for determining the ongoing conservation status of these salamanders, for guiding their management, and for furthering our understanding of how organisms utilize karst habitats.

## Material and Methods

### DNA sequencing and data collection

Sampling for this investigation included extensive collections of the focal *E. neotenes* complex (including the nominal taxa *E. pterophila*, *E. neotenes* and *E. latitans*). We also included samples of *E. nana*, which is closely related to the *E. neotenes* complex (Bendik *et al*., 2013; Chippindale *et al*., 2000; Devitt *et al*., 2019) and samples from further west of the focal area for comparison. These western samples are historically assignable to *E. troglodytes* and “*E.* sp. 2” following Devitt *et al*. (2019) who recognized significant differentiation within *E. troglodytes* (Table 1). Tissues from tail clips were collected from 374 wild-caught individual salamanders from 2019 to 2021. Sampling was augmented with archived material from three sources: 17 tissue samples from museum specimens housed at the Texas Natural History Collections, Texas Memorial Museum at The University of Texas at Austin, Austin, Texas, collected from 1990-1994, DNA samples from 10 individuals from Preserve Cave donated by Tom Devitt, and DNA samples from 242 salamanders generated for a previous study from individuals collected from 2004 and 2005 (Lucas *et al*., 2009) (Table 1, Figure 1). All tissue samples were preserved in 95% EtOH. Samples from Lucas *et al*. (2009) were extracted using the Promega Wizard SV Genomic DNA Purification System and the Gentra Systems Puregene DNA Purification kits. DNA from all other samples was extracted using the DNeasy Blood and Tissue Kit (Qiagen Inc., Alameda, CA, USA).

Following the methods of Parchman *et al*. (2012) and Gompert *et al*. (2014), we constructed a reduced representation genomic library for each salamander. For these libraries, genomic DNA was digested with the EcoR1 and Mse1 restriction enzymes. To the resulting fragments, Illumina adaptors with unique 8-10bp individual multiplex identifier (MID) sequences were ligated. Fragments were then amplified with two rounds of PCR using iProof high fidelity polymerase (BioRad, Inc.). Amplicons from these PCR reactions were then pooled. Fragments between 300 - 450bp were selected using a BluePippin (Sage Science Inc., Beverly, MA, USA) and the resulting fragments were sequenced on two lanes of Illumina Novaseq 6000 (SR 100) at the University of Texas Austin Genomic Sequencing and Analysis Facility (Austin, Texas, USA).

### Assembly and variant calling

From the resulting sequence reads, PhiX reads were removed by assembly to the PhiX genome using bowtie version 1.1.2 (Langmead *et al*., 2009). Custom scripts were used to remove MIDs from each read and to filter short reads and reads that contained Mse1 adapter sequence. The resulting sequence reads were written to individual files in fastq format (median = 1,042,576 sequence reads per individual). Thirty-six individuals each produced fewer than 400,000 reads and were excluded from further analyses. Excluded individuals originated from 21 distinct localities and included a maximum of four individuals from any one locality. One specimen from the Texas Natural History Collections (from Pfeiffer Ranch (16); we refer to specific localities by name and corresponding number following Table 1 in parentheses) was among the excluded individuals. The reads from the remaining 607 individuals (Table 1, Figure 1) were filtered and clustered using the strategy described in the dDocent variant calling pipeline (Puritz *et al*., 2014) and CD-hit (Fu *et al*., 2012). We ignored sequence reads with less than four copies per individual and shared among less than four individuals. The filtered reads were assembled with a homology threshold of 80% (other thresholds from 80-95% were investigated but produced very similar assemblies, data not presented). The consensus reads from this *de novo* assembly were used as the basis for a reference-based assembly of all reads using the *aln* and *samse* algorithms of the Burrows-Wheeler Aligner (bwa version: 0.7.12) (Li & Durbin, 2009) with a seed length of 20 and up to four mismatches allowed.

Following assembly, we identified variable sites, or polymorphic single nucleotide sites (SNPs), with bcftools version 1.9 (Li *et al*., 2009) using the *mpileup* and *call* commands, ignoring indels and retaining variable sites if the posterior probability that the nucleotide was invariant was *<*0.05. We performed additional filtering on the Variant Call Format file using custom scripts to exclude variable sites with sequence depth less than 607 reads (an average of one read per site per individual, 1X) and greater than 47,246 reads (equal to the mean sequence depth across sites plus two standard deviations; this filter is designed to remove potentially paralogous reads), less than at least 20 reads of the alternative allele, mapping quality less than 30, an absolute value of the mapping quality rank sum test greater than 1.96, an absolute value of the read position rank sum test greater than 1.96, an absolute value of the base quality rank sum test greater than 1.96, minor allele frequency less than 0.05, or missing data for more than 303 individuals (50% of individuals). One variable site per contig was chosen randomly and retained to maximize independence among loci.

### Analyses

To answer our first two questions pertaining to the organization of genetic variation, we estimated genotypes, allele frequencies and admixture proportions and used these to illustrate patterns of genomic variation. We estimated admixture proportions for each individual, allele frequencies for each cluster (population) and posterior genotype probabilities for all loci for all individuals using the Bayesian admixture algorithm entropy (Gompert *et al*., 2014; Shastry *et al*., 2021). A series of models were fit to the data varying the number of clusters or populations (k) from two to 15. Two MCMC simulations (chains) of 105,000 steps, with a burnin of 5,000 steps, retaining every 10th step (total of 10,000 steps for each chain), were run for each k. We calculated Gelman and Rubin’s convergence diagnostic and effective sample sizes (Brooks & Gelman, 1998; Gelman & Rubin, 1992) for each chain with the package coda version 0.19-1 in R (Plummer *et al*., 2006; R Core Team, 2022) to assess model performance and check that the models reached a stable sampling distribution. We rejected model results when the mean Gelman and Rubin’s convergence diagnostic across individual values of the admixture proportion, *q*, was greater than 1.1 or when increasing k did not result in a recognizable cluster. Models for k=2 - 5 satisfied these criteria. Attempts to employ more MCMC steps, longer burnin, and different thinning strategies failed to rescue models with higher numbers of clusters (data not presented). Posterior estimates of genotypes were then averaged across all MCMC steps for models k = 2 - 5. Patterns of variation among individuals were illustrated by ordination of multilocus genotypes using Principal Component Analysis (PCA), and patterns of admixture were illustrated with barplots, all performed in R (R Core Team, 2022). To quantify differentiation among sites, we calculated genome-average Nei’s *G_ST_* (an analog of the standard measure of differentiation, 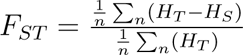) for all pairwise combinations of sites, which we refer to as *F_ST_*. *F_ST_* was calculated as the average across all loci and 1000 bootstrap resamples were used to calculate 95% confidence intervals.

We then used samtools version 0.1.9 to calculate two measures of genetic diversity for all aligned nucleotide sites for each of the 29 localities with samples of *≥* 8 individuals. We estimated Watterson’s *θ* (based on the number of segregating sites) and Tajima’s *π* (nucleotide diversity or heterozygosity) using the expectation-maximization algorithm with 20 iterations (Li, 2011), which was sufficient for values of both metrics to converge for all localities.

To specifically address our first question regarding whether genomic variation is organized according to the nominal taxonomy (i.e. three nominal species in the *E. neotenes* complex), we employed two approaches: variance partitioning using Redundancy Analysis (RDA) (Capblancq & Forester, 2021) and a Bayesian linear mixed modeling approach (Gompert *et al*., 2014). First, we used RDA to partition genetic variance using predictors representing the historical nominal taxonomy plus features of geology (aquifer zones) as a comparison, and geographical space. These analyses were performed in R (R Core Team, 2022). To the best of our ability, we assigned localities to a nominal species based on historical descriptions and previous taxonomic studies that employed molecular marker data (Bendik *et al*., 2013; Chippindale *et al*., 2000; Devitt *et al*., 2019). It should be noted that because many of our sampling localities have not been sampled previously, there is a substantial possibility for error in these assignments. These nominal species designations (Table 1) were used as predictors of genomic variation in RDAs. Similarly, we used major aquifer designation for each locality as predictors (Table 1). The localities included sites located in the Edwards - Trinity Aquifer in the western portion of the Edwards Plateau, sites in the Trinity Aquifer in the northern regions of our sampling area, and sites in the Balcones Fault Zone of the Edwards Aquifer along the southeastern edge of our sampling area (Table 1, Figure 1). We accounted for geography by using Moran’s Eigenvector Mapping functions (MEMs) which describe spatial autocorrelation of our sampling sites at multiple scales (Borcard & Legendre, 2002; Borcard *et al*., 2004; Dray *et al*., 2006; Forester *et al*., 2018). MEMs were calculated using the *dbmem* function from the adespatial package. We converted the posterior genotype matrix (described above) into a Euclidean distance matrix among individuals using the *vegdist* function in the vegan package. (This does not alter results of any of the analyses but does reduce computational time.) We performed RDA using nominal taxonomy, aquifer, and MEMs as predictors of genomic variation using the *dbrda* function from the vegan package followed by permutational ANOVA. We partitioned variance for each of the predictors using the *varpart* function in the vegan package.

In a second method for addressing whether genetic variation was organized along nominal taxonomic divisions or major aquifers, we constructed a Bayesian mixed model regression to compare geographic distance, “taxonomic” distance and “aquifer” distance as predictors of population differentiation (Gompert *et al*., 2014). Specifically, we used this model to ask whether population differentiation was best explained by geography, nominal taxonomy, major aquifer or combinations of these predictors. The regression model accounts for the correlated error structure in pairwise distance measures (genetic, geographic, taxonomic and aquifer) by incorporating random effects for each population that represent mean population specific deviations in the distance matrices following Clarke *et al*. (2002). Thus,

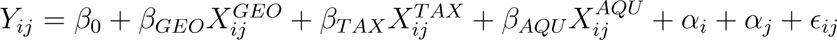

where *Y_ij_* is pairwise genetic distance between populations *i* and *j*, 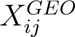 is the corresponding geographic distance, 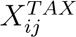 indicates taxonomic identity (with 1 when populations *i* and *j* are from different nominal taxa and 0 when from the same nominal taxon), and 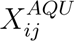 indicates major aquifer identity (with 1 when populations *i* and *j* are located in different major aquifers and 0 when from the same major aquifer) (Table 1). The *β*’s are fixed effect regression coefficients. The *α* terms represent random effects for individual populations (Clarke *et al*., 2002) and *ɛ_ij_* is the error. For genetic distances, we used pairwise *F_ST_ /*(1 *−F_ST_*) (Rousset, 1997), (*F_ST_* calculated as described above and for all localities with *n ≥* 8). Pairwise genetic distances were log transformed and centered. The geographic distance matrix comprised great circle distances between localities expressed in meters which were log transformed, centered and scaled. We used uninformative Gaussian priors for the regression coefficients (*β*’s) (*µ* = 0*, σ*^2^ = 1000) and hierarchical Gaussian priors for the population random effects (*α*’s) (*µ* = 0*, σ*^2^ = 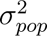) and uninformative gamma priors for the random effects and residual variances (as reciprocals) (*α* = 1*, β* = 0.01). Models were implemented in r using rjags with MCMC run in jags (Plummer *et al*., 2003). All models included three MCMC chains of 10,000 iterations, a burnin of 2000 iterations and thinning every five steps. Models were constructed for the entire data set of localities with *n ≥* 8 (29 localities) and for a “focal” data set of localities assigned to the *E. neotenes* complex (23 localities). We fit full models with terms for geography, taxonomy and aquifer, plus models with two-way combinations of predictors, and models with single predictors, and used Deviance Information Criterion (DIC) for model comparison.

Our third question regarding the possibility of gene exchange during the history of these populations presents challenges because observed admixture could result from incomplete lineage sorting (ancestral polymorphism), historic gene flow, contemporary gene flow, or combinations of these (e.g. Holder *et al*., 2001; Joly *et al*., 2009; Machado *et al*., 2002; Sang & Zhong, 2000). It should be noted that many methods for inference of gene flow, including those used here, cannot readily discriminate between secondary contact with gene flow versus gene flow during divergence (Endler, 1977; Pinho & Hey, 2010).

Given these difficulties, we employed three analyses with different approaches in an attempt to understand the history of gene exchange: We first used the methods of Pickrell & Pritchard (2012) as implemented in treemix version 1.12. We used allele frequencies from all focal sampling localities with sample sizes of at least eight individuals to estimate the hypothesized population graph representing the evolutionary history of the sampled populations assuming no migration or admixture (*m* = 0), with the sample from Fessenden Springs (2) designated as the outgroup. We then sequentially evaluated the contribution of additional migration nodes (admixture events, *m* = 1-10). Each model of *m* migration nodes was replicated with 1000 runs and the proportion of the total explained variance in allele frequencies was calculated for each replicate using a custom Rscript written by D. Card (https://github.com/darencard/RADpipe/blob/master/treemixVarianceExplained.R). The asymptote in the median proportion of variance explained across replicates for each *m* was used as a guide to determine whether adding migration events improved model fit. Population graphs for *m* = 0 and for *m* with the highest median variance explained were plotted using the R package ggtree (Yu *et al*., 2017).

In a second analysis of gene exchange, we calculated *f*_3_ statistics (Patterson *et al*., 2012; Reich *et al*., 2009) with the threepop program from treemix version 1.12. *f*_3_ statistics specifically test the hypothesis that the evolution of sampled populations conforms to the expectations of a bifurcating model (i.e. the null hypothesis of no history of gene exchange). The test is conducted for sets of three populations, two potential source populations and a target population. Significantly negative *f*_3_ statistics indicate a departure from a strictly bifurcating evolutionary history for the three populations due to population admixture in the target population. We calculated *f*_3_ statistics for all sampling localities included in the treemix analyses described above.

As a complement to the above analyses designed to detect gene flow, we used EEMS (version 0.0.0.9000) (Petkova *et al*., 2016) to estimate an effective migration surface which provides a picture of the geography of departures from patterns of isolation-by-distance (IBD). This effective migration surface identifies areas with low or high gene flow relative to a stepping-stone model. Our expectation is that gene flow among localities and lineages of *Eurycea* salamanders should appear as departures from the IBD expectation such that the effective migration surface will illustrate the presence of barriers to and/or corridors of gene flow. If, on the other hand, the history of divergence in these salamanders is one of primary divergence with little or no gene flow, we expect to observe few or no departures from IBD expectations. We estimated the effective migration surface assuming 700 demes and used three MCMC chains with 6 million iterations each, a burnin of 3 million iterations and a thinning interval or 10,000 iterations.

## Results

Sequencing produced 671,939,934 sequence reads with MIDs for 607 individuals (median = 1,072,524 reads per individual). 42,274,123 reads were retained in the final assembly after filtering from which a SNP set 16,094 loci was called. Mean sequence depth was 4.3 reads per individual per locus with an average of 26.4% missing data per locus. Ordination of posterior genotype estimates from entropy showed differentiation of the western localities, *E. troglodytes* and *E.* sp.2 (*sensu* Devitt *et al*., 2019) from all other samples on PC axis 1 which explained about 11% of the total genomic variation (Figure 2). Samples of *E. nana* were also differentiated from the focal *E. neotenes* complex, but clearly closely related. The focal *E. neotenes* complex separated into three less differentiated clusters on PC axis 2 which explained less than 5% of the variance. One group of individuals from localities in the southwestern portion of our sampling (localities 10 - 14, 17, 19, 21, 22, Table 1, Figure 1) formed a distinct cluster (colored variously orange in Figure 2). The other localities north (localities 5 - 9, 15, 16, 18, 20, 23 - 27 and colored green in Figure 2) and east (including 28 - 35 and colored purple/magenta in Figure 2) of the southwestern (orange) cluster are less differentiated from each other.

**Figure 2:**
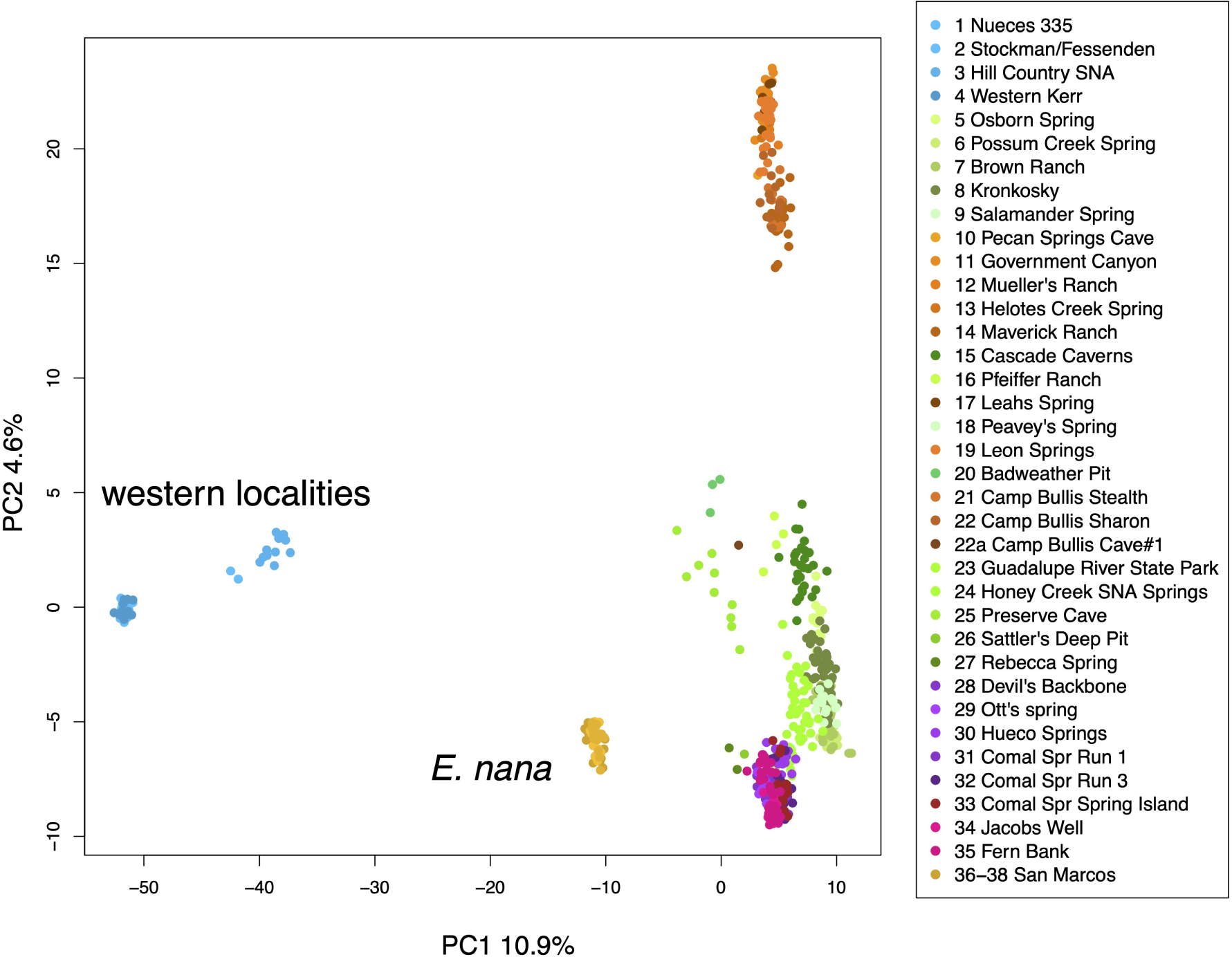
Principal Components Analysis of posterior genotype estimates summarizing variation among 607 *Eurycea* individuals based on 16,094 SNP loci. Each individual is represented by a point colored by locality (Table 1, Figure 1).

Clustering analyses for k *>* 5 failed to meet our criteria for convergence and we focus on results of k = 2 to 5. DIC scores were lowest for MCMC chains at k = 5 (Supplementary Figure 1). Barplots of admixture proportions (Figure 3) mirror the apparent divisions in the PCA (Figure 2). The western localities (1 - 4) are differentiated first, followed by *E. nana* (36 - 38) at k = 4. At k = 5, the focal *E. neotenes* complex comprises three clusters as in the PCA. However, the clusters are not distinct and many localities show signs of admixture (Figures 3, 4). Admixture is particularly evident at Cascade Caverns (15), the type locality for *E. latitans* (Smith & Potter, 1946), Guadalupe River State Park (23), Hueco Springs (30), and all three Comal Springs sites (31-33). Honey Creek SNA (24), shows ancestry from all three groups within the *E. neotenes* complex, while Sattler’s Deep Pit (26) and Rebecca Spring (27) show variable ancestry from 4 groups: from two of the *E. neotenes* complex (green and purple groups), from *E. nana*, and also from the western localities. Preserve Cave (25) shows ancestry from all five clusters. Further evidence of admixture is seen in the western localities, particularly Nueces 335 (1) and Hill Country State Natural Area (3) which share ancestry with the orange group and *E. nana*. *E. nana* ancestry is also observed in some individuals from Jacobs Well (35) and Fern Bank (35).

**Figure 3:**
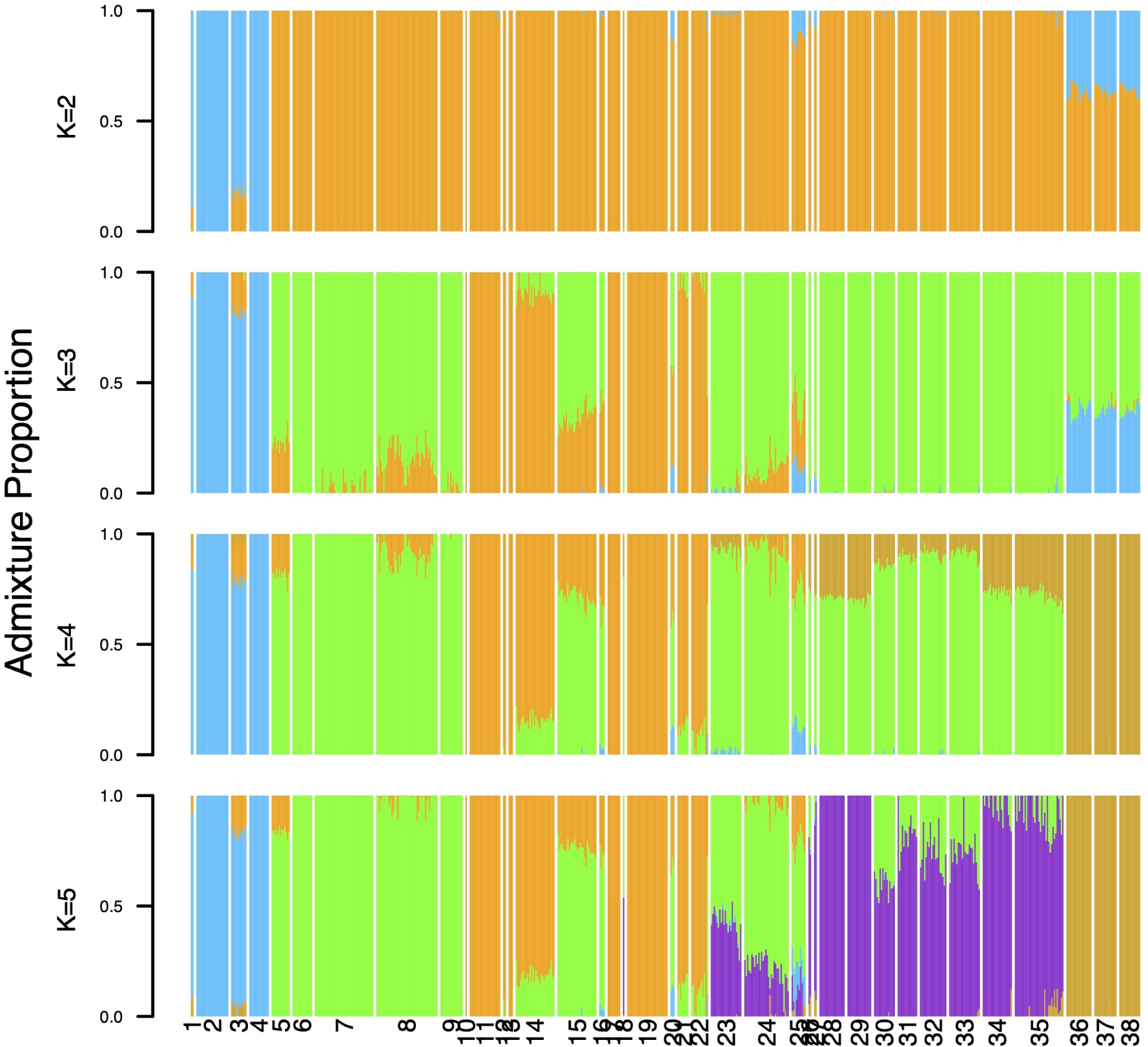
Admixture proportions for k = 2 - 5. Localities are oriented by longitude (west to east, Figure 1, Figure 4). Locality numbers follow Table 1.

**Figure 4:**
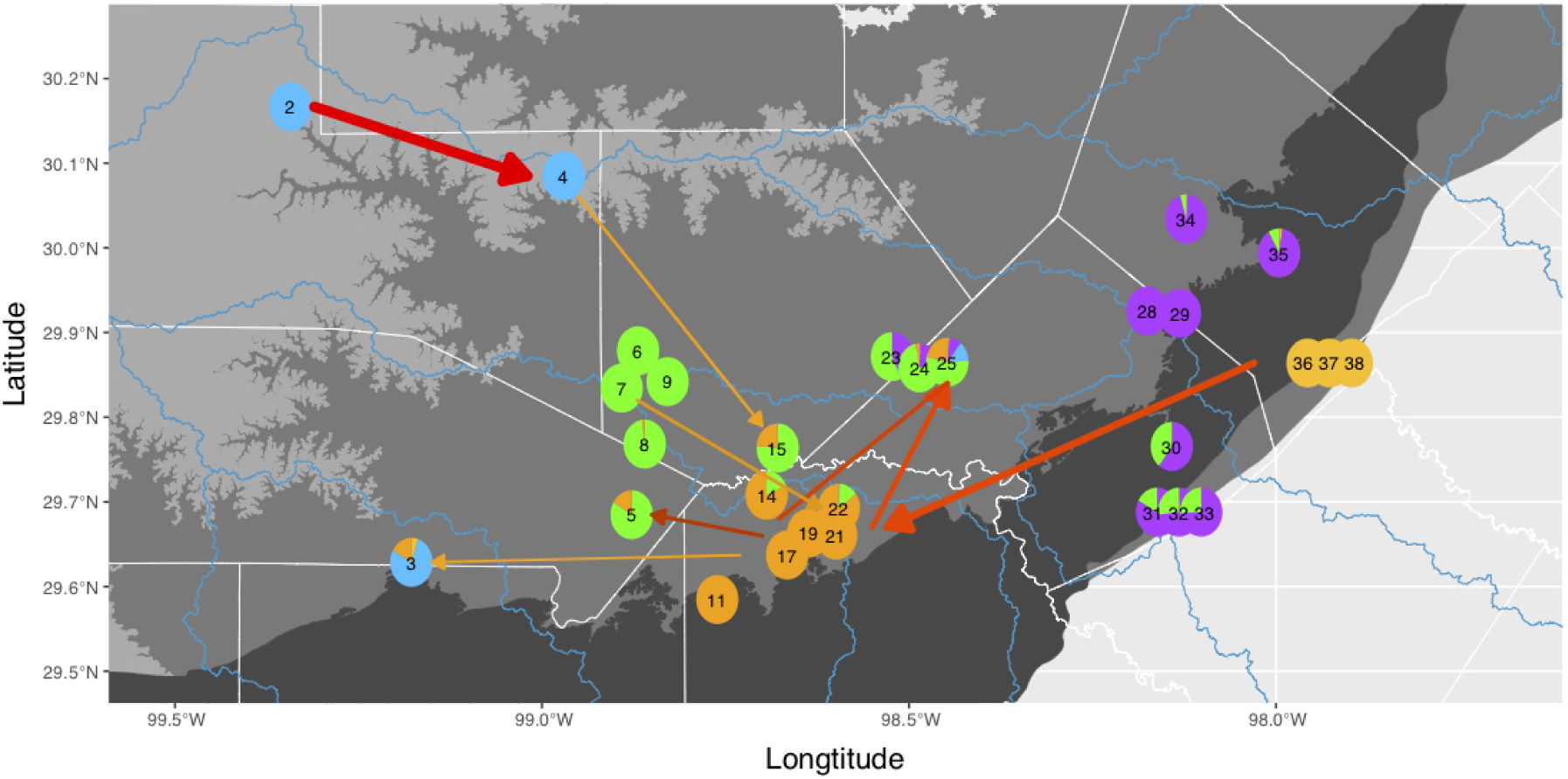
Amixture proportions per locality and migration events inferred from treemix. Pie diagrams represent proportional ancestry (q) for localities with at least eight sampled individuals based on entropy clustering for k=5. Arrows show the direction of inferred migration events with color and magnitude as in Figure 9. Note that inferred migration events between localities cannot be interpreted as the actual paths of gene flow that occurred in the aquifers. Locality numbers follow Table 1.

Overall, levels of differentiation are low, especially within the *E. neotenes* complex. Pairwise *F_ST_*‘s are very low (Supplementary Tables 1-4). The largest observed value (*F_ST_* = 0.126, 95% confidence interval: 0.122-0.131) was between Western Kerr (4) and Leahs Springs (17). Excluding the western populations and *E. nana* (i.e. within the focal *E. neotenes* group), *F_ST_*’s ranged from 0.004 (0.004 - 0.004) (between Brown Ranch (7) and Albert and Bessie Kronkosky State Natural Area (8)) to 0.04 (0.038 - 0.042) between Leahs Springs (17) and Devil’s Backbone (28), and 0.04 (0.039 - 0.042) between Leahs Springs (17) and Ott’s Springs (29)). These results are similar to those reported by Devitt *et al*. (2019) for this group (eastern *Blepsimogle*) and this level of differentiation is of the same magnitude as, or slightly higher than, population differentiation observed within another central Texas *Eurcyea*, *E. chisholmensis* (Nice *et al*., 2021).

Genetic diversity was slightly lower in the *E. neotenes* complex localities compared to *E. nana* or the western localities, especially for Watterson’s *θ* (Figure 5). Diversity across all sites was roughly a third to a half of the diversity measured by similar methods in *E. chisholmensis* (Nice *et al*., 2021). However, there were no obvious disparities in diversity across the *E. neotenes* complex.

**Figure 5:**
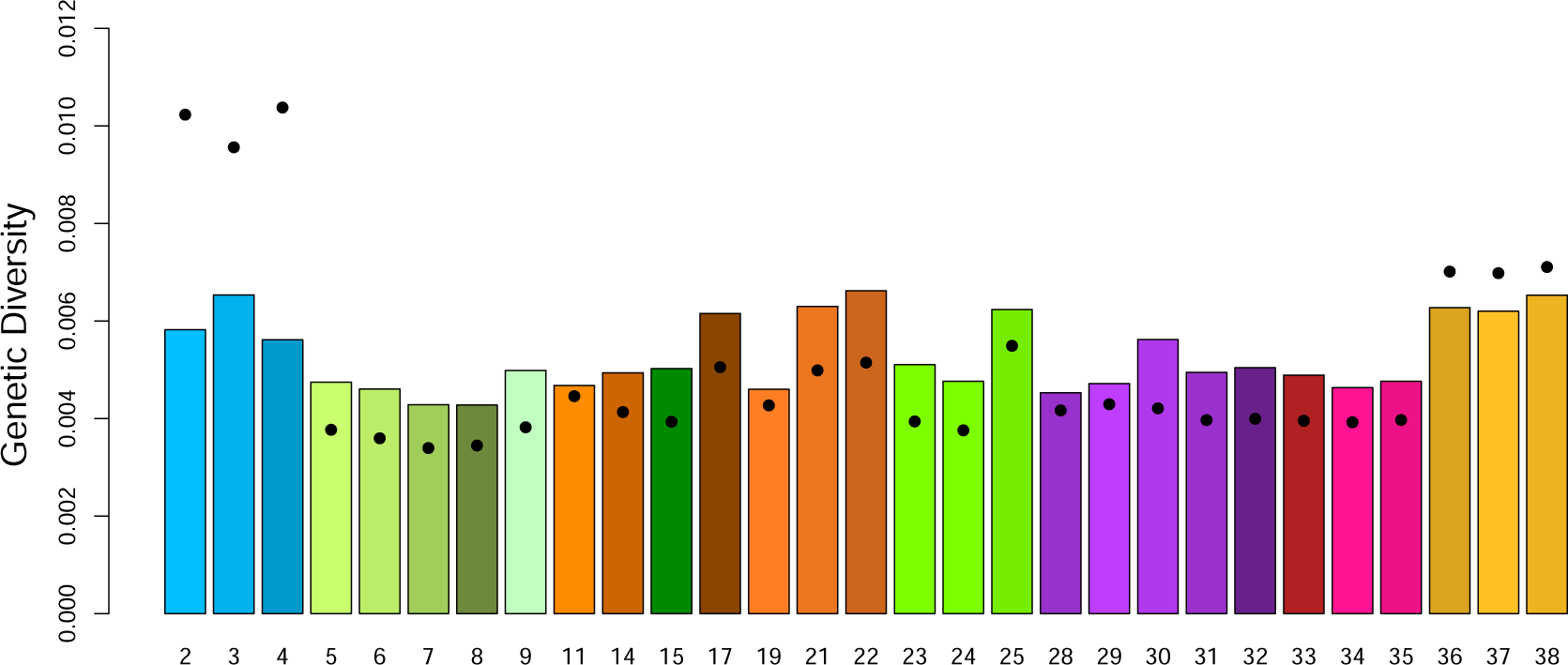
Genomic diversity estimated for *Eurycea* salamanders for all aligned sites from localities with at least eight sampled individuals. Nucleotide diversity as Tajima’s *π* (expected heterozygosity) is plotted as bars. Watterson’s *θ* is indicated as points for each site. Localities are arranged from west to east and locality numbers follow Table 1 and Figure 1.

Predicting genotypic variation by nominal taxonomy and major aquifer (Table 1) and spatial autocorrelation using RDA revealed both low levels of variance explained and collinearity among predictors. In initial models, taxonomy alone (i.e., not controlling for spatial variation) explained 17.0% of total genotypic variation, while major aquifer alone explained 9.7%. Space, as captured by eight MEMs, all of which were significantly associated with genotypic variation, explained 13.3%. However, the combined model with nominal taxonomy, major aquifer and eight MEMs explained only 21% of the variance. In this combined model, taxonomy explained 4.7%, aquifer explained less than 1% and space accounted for 3.5% of the total genotypic variation (Figure 6), but the overlap of all three predictors (i.e. the variance explained by the combination of all three) was 6.3%, and two-way entanglements between the predictors accounted for the remaining variance explained (Figure 6). In summary, spatial variation in the distribution of genetic variation in these populations is collinear with taxonomy and the distribution of aquifers (Devitt *et al*., 2019).

**Figure 6:**
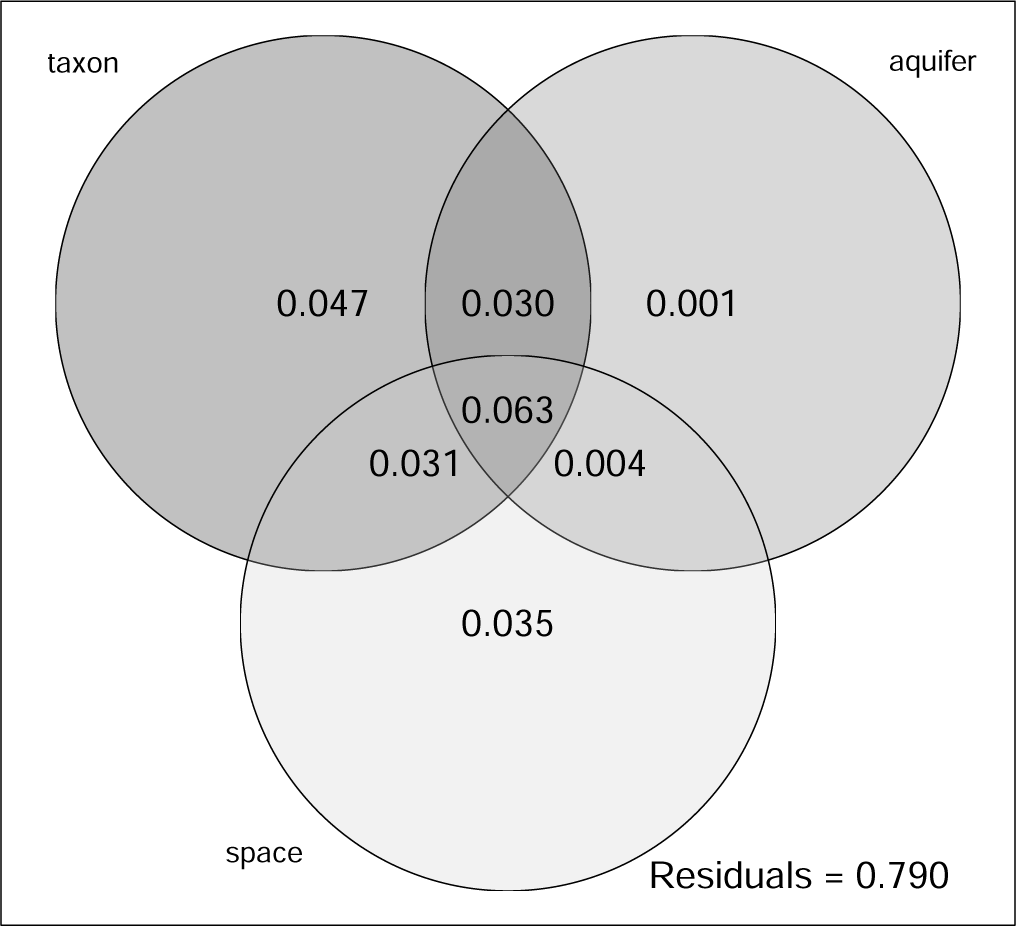
Venn diagram showing results of variance partitioning of the total genetic variance into components representing nominal taxonomy (taxon), major aquifer association (aquifer), and spatial autocorrelation (space) as captured by Moran’s eigenvector mapping functions (MEMs). The full model explained 21% of the total variance.

The geographical signal detected in the examination of patterns of differentiation using RDA is evident when examining the relationship between pairwise differentiation and geographic distance (Figure 7). Within the three nominal focal species, differentiation increases mostly linearly with distance between localities (Figure 7, Supplementary Figure 2). The Bayesian regression showed that more distant localities, localities classified as different taxa, and localities in different major aquifers were more genetically distinct (Table 3). Based on DIC, a model with taxonomy and geography and the the full model (with terms for taxonomy, geography and aquifer) received the most support for all localities and for just the focal *E. neotenes* complex localities. However, reduced models that included geographic distance, including the model with just geographic distances, were preferred over alternate reduced models that did not include geographic distances for all localities and for the focal localities (Table 3). Thus, as found with the RDA, genetic variation appears to be weakly structured along both geographic and taxonomic dimensions, and to a lesser extent, major aquifer, all of which are collinear.

**Figure 7:**
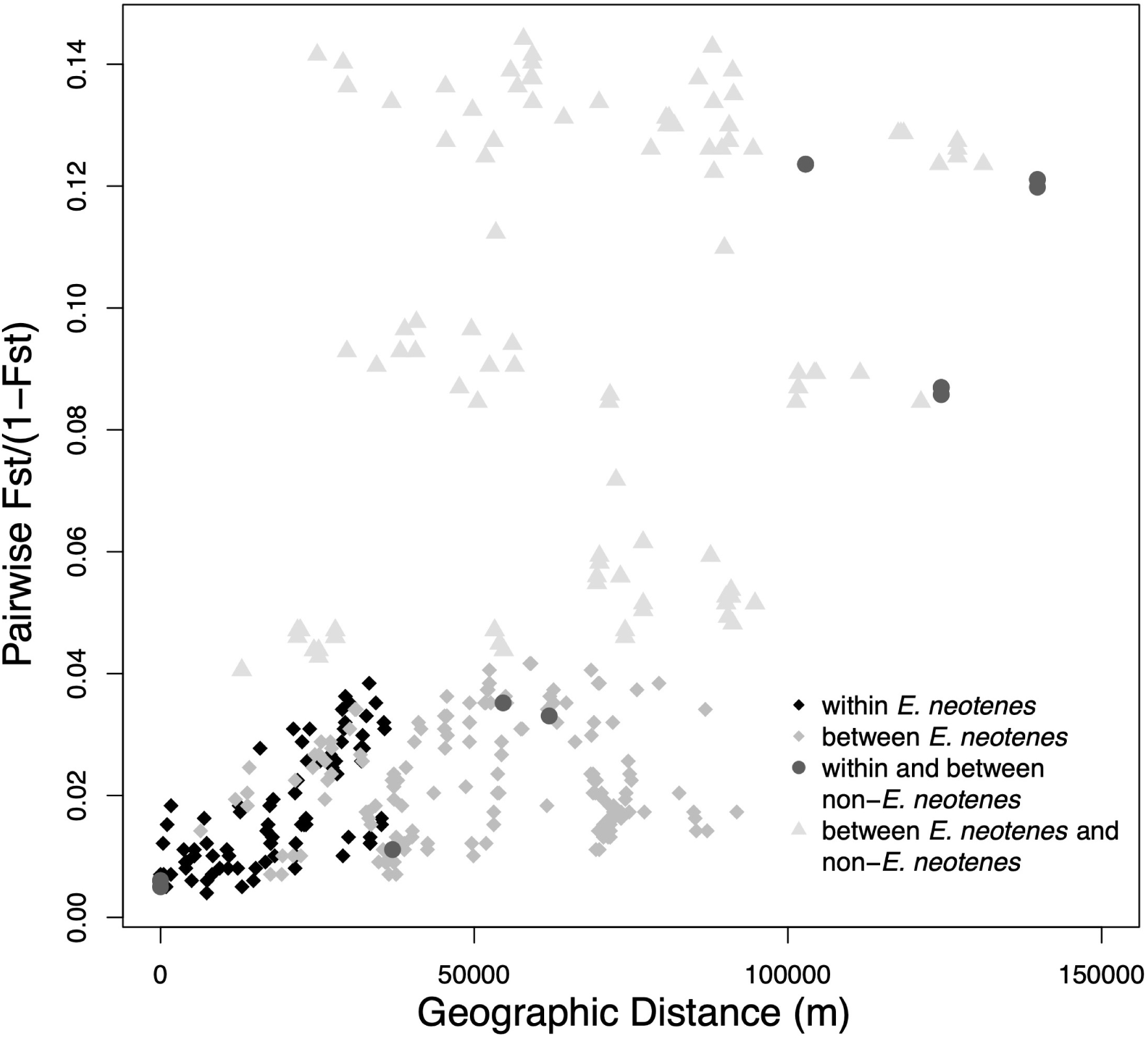
The relationship between geographical distance and genetic distance between localities as measured by *F_ST_*. Symbols and shading indicate different types of pairwise comparisons, including within the *E. neotenes* complex (i.e., comparisons between localities within each of the three nominal taxa), between the *E. neotenes* complex taxa (e.g. between *E. neotenes* and *E. pterophila*), within and between non-*E. neotenes* taxa (e.g. within and between *E. troglodytes* and *E. nana*) and between *E. neotenes* complex taxa and non-*E. neotenes* taxa (e.g. between *E. latitans* and *E. nana*).

TREEMIX analyses provided some evidence of gene exchange among populations. Adding migration events continued to improve the amount of variance explained until reaching an asymptote around eight migration events (Figure 8). The initial drift tree (population graph with no migration events, m = 0) has patterns similar to the PCA and barplots (Figure 9A) and explained considerable variance in allele frequencies. Adding migration events improved model fit though the increase in variance explained was small (Figure 8). The population graph incorporating eight migration events (m = 8) (Figure 9B) shows numerous migration events among *E. neotenes* complex populations, commonly involving the southwestern localities (localities 11, 14, 17, 19, 21, 22). There is also evidence of historical gene exchange between the western localities (*E. troglodytes* and *E.* sp. 2) and *E. neotenes* complex populations and an historical gene flow event from *E. nana* to the southwestern localities. Many of these migration events parallel the patterns of admixture in the clustering analyses (Figure 3). Interestingly, the direction of these migration events indicates multiple paths for gene flow (Figure 4) and that gene flow appears to have been relatively unrestricted in geographical space. Supplementary Figures 3 and 4 present population graphs with seven (m = 7) and nine (m = 9) migration events.

**Figure 8:**
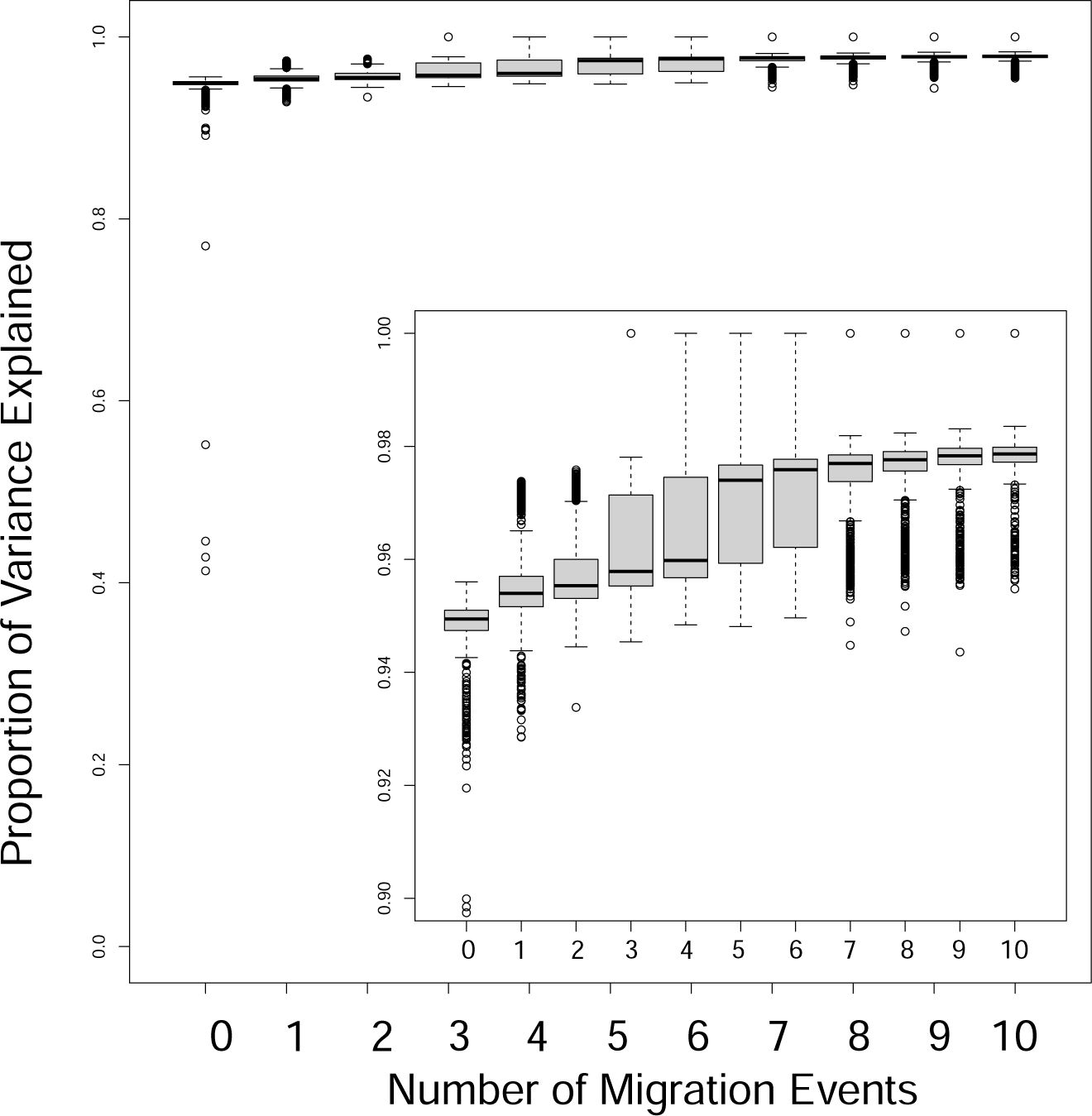
Boxplots of variance explained from Treemix analyses of salamander localities with 0 - 10 migration events (m = 0 - 10). Boxplots show mean variance explained and variation from 1000 bootstrap replicates for each number of migration events, *m*. Inset shows magnified boxplots.

**Figure 9:**
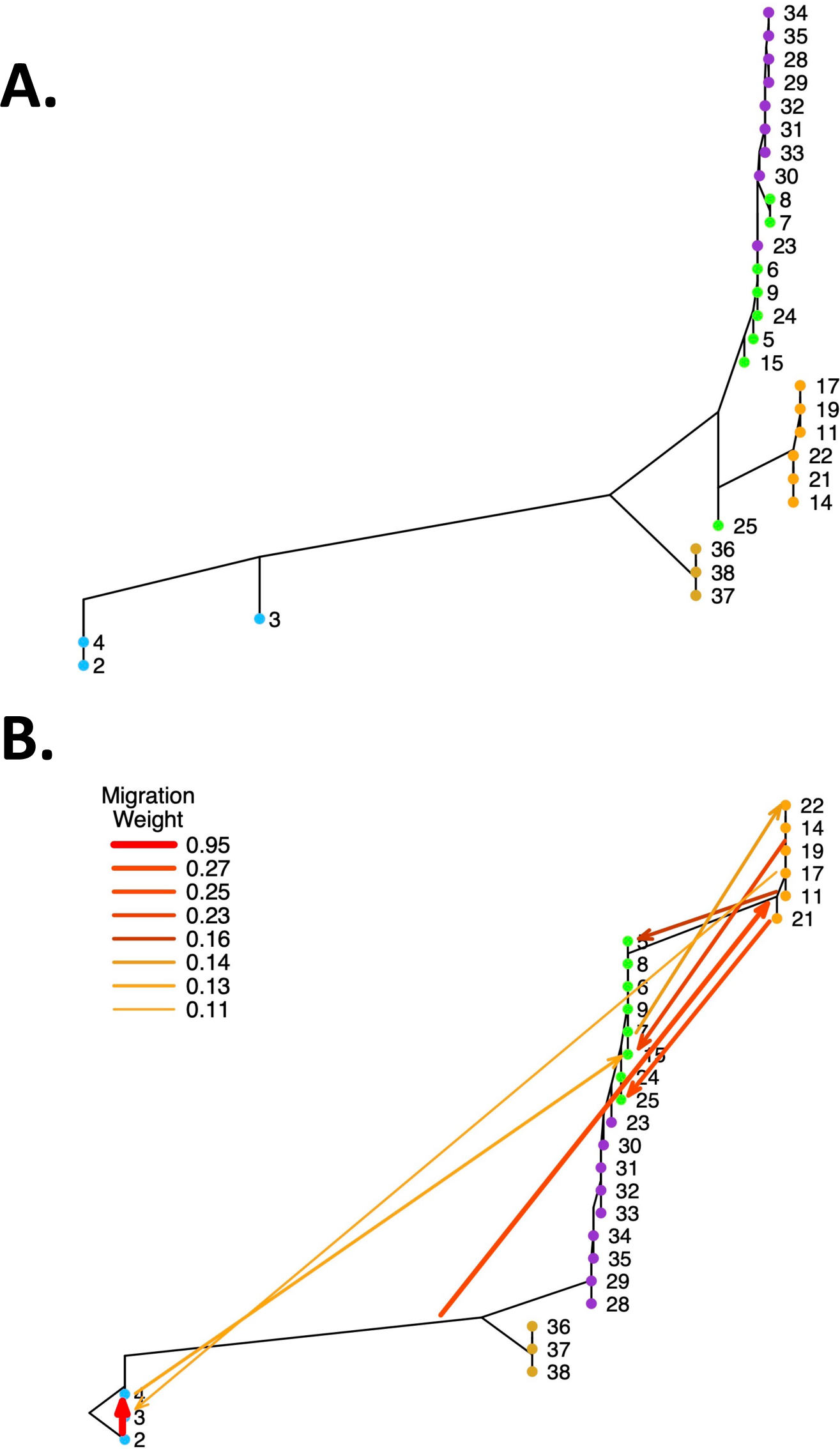
Treemix results. A) Drift tree (population graph with no migration, *m* = 0) of focal salamander localities with minimum sample sizes of at least eight individuals. Locality numbers follow Table 1. The Fessenden Springs site *E.* sp. 2, locality 2, serves as the outgroup. B) Treemix population graph of focal salamander localities with eight migration events (m = 8). Locality numbers follow Table 1.

Significantly negative *f*_3_ statistics were detected in more than 48% of the 10,962 three-locality comparisons and included every locality as a target (i.e. receiving gene flow from the source populations) (Table 2). These results indicate that the history of the central Texas *Eurycea* is inconsistent with a bifurcating pattern of divergence and are an indication of a history of some gene exchange among localities. These results, in combination with the treemix results, suggest that admixture among localities has been a component of the history of these salamanders.

**Table 2:** Results of three-population test of admixture. These are abbreviated results with only the most negative *f*_3_ statistic for each locality as the target of admixture presented along with the source localities for that specific test. For localities 31-33 and 36-38, localities within large spring complexes (Comal Springs and San Marcos Springs, respectively), the most negative *f*_3_ statistics were observed with source localities located in the same spring complex. Below the line we report the second-most negative *f*_3_ statistics for these sites (i.e. with one source population from outside the local spring complex), and the most negative *f*_3_ statistic in which both source populations are outside of the local spring complex. Tests in bold are significant at an arbitrary p *<* 0.000001. Only localities with n *≥* 8 were included in these tests.

| Target | Source 1 | Source 2 | $f_3$ | Std. Error | Z-score |
| --- | --- | --- | --- | --- | --- |
| 2 | 3 | 4 | -0.00229295 | 0.000106912 | -21.4471 |
| 3 | 2 | 19 | -0.00957279 | 0.000277759 | -34.4644 |
| 4 | 2 | 3 | -0.00353063 | 0.000114193 | -30.918 |
| 5 | 8 | 15 | -0.00636996 | 7.62176e-05 | -83.576 |
| 6 | 7 | 24 | -0.00638172 | 5.60728e-05 | -113.811 |
| 7 | 8 | 9 | -0.00236467 | 2.95913e-05 | -79.9111 |
| 8 | 5 | 7 | -0.00225418 | 4.02737e-05 | -55.9714 |
| 9 | 7 | 8 | -0.0055665 | 4.44625e-05 | -125.195 |
| 11 | 14 | 19 | -0.00355891 | 5.39144e-05 | -66.0104 |
| 14 | 15 | 19 | -0.00452096 | 6.79268e-05 | -66.5564 |
| 15 | 8 | 25 | -0.0040262 | 7.09935e-05 | -56.7122 |
| 17 | 14 | 19 | -0.00919705 | 7.65707e-05 | -120.112 |
| 19 | 11 | 14 | -0.00349781 | 3.71554e-05 | -94.1402 |
| 21 | 14 | 22 | -0.010668 | 9.59334e-05 | -111.202 |
| 22 | 14 | 21 | -0.00868404 | 7.10733e-05 | -122.184 |
| 23 | 24 | 30 | -0.00410975 | 5.4395e-05 | -75.5538 |
| 24 | 15 | 23 | -0.00286774 | 5.34846e-05 | -53.6182 |
| 25 | 3 | 15 | -0.0106783 | 0.000196798 | -54.2601 |
| 28 | 29 | 30 | -0.00506417 | 6.39127e-05 | -79.2358 |
| 29 | 28 | 30 | -0.00563487 | 5.82594e-05 | -96.7203 |
| 30 | 24 | 35 | -0.00610045 | 6.66204e-05 | -91.5703 |
| 31 | 32 | 33 | -0.00663148 | 5.03875e-05 | -131.61 |
| 32 | 31 | 33 | -0.00536484 | 4.2545e-05 | -126.098 |
| 33 | 31 | 32 | -0.00411446 | 4.88667e-05 | -84.1977 |
| 34 | 23 | 35 | -0.00368665 | 7.33839e-05 | -50.2379 |
| 35 | 23 | 34 | -0.00304483 | 5.48816e-05 | -55.48 |
| 36 | 37 | 38 | -0.00484558 | 3.77361e-05 | -128.407 |
| 37 | 36 | 38 | -0.00545892 | 3.82996e-05 | -142.532 |
| 38 | 36 | 37 | -0.00573825 | 4.21412e-05 | -136.167 |
| 31 | 32 | 35 | -0.00659928 | 6.00073e-05 | -109.975 |
| 31 | 23 | 29 | -0.00538465 | 1.23327e-04 | -43.6617 |
| 32 | 24 | 31 | -0.00563032 | 5.22206e-05 | -107.818 |
| 32 | 7 | 29 | -0.00446188 | 1.23057e-04 | -36.25880 |
| 33 | 30 | 32 | -0.00431445 | 5.20231e-05 | -82.9335 |
| 33 | 7 | 29 | -0.003547700 | 1.29704e-04 | -27.35230 |
| 36 | 35 | 38 | -0.00490134 | 6.40483e-05 | -76.5256 |
| 36 | 34 | 6 | 0.00232263 | 4.79394e-04 | 4.84493 |
| 37 | 35 | 36 | -0.00529426 | 6.64943e-05 | -79.61980 |
| 37 | 2 | 9 | 0.00211488 | 4.70187e-04 | 4.49795 |
| 38 | 35 | 36 | -0.00568249 | 6.85136e-05 | -82.93960 |
| 38 | 2 | 9 | 0.00169419 | 4.70917e-04 | 3.59764 |

The estimated effective migration surface showed distinct areas where gene flow was higher or lower than expected under a two-dimensional stepping-stone model (Figure 10). A corridor (an area of higher gene flow (*m*), or lower differentiation than expected given the geographic distances) was observed connecting the eastern localities at Devil’s Backbone (28) and Ott’s Spring (29) with the Comal Springs complex localities (sites 30 - 33) to the south, Guadalupe River State Park (23), Honey Creek (24) and Preserve Cave (25) in the northern portion of sampling, and to south westerly localities including Camp Bullis (21,22). Another corridor connects the western localities of Fessenden Springs (2), Hill Country SNA (3) and Western Kerr (4) with southwestern sites Osborn Springs (5), Albert and Bessie Kronkowsky SNA (8) and Government Canyon (11). Another smaller corridor connects Osborn Springs (5) with Maverick Ranch (14) and Cascade Caverns (15). These areas of higher inferred gene flow roughly parallel the inferred migration paths from Treemix (Figure 9) and the patterns of admixture (Figure 3). There are also several areas that appear as barriers (or areas with lower than expected gene flow or higher differentiation than expected given the geographic distances). One major barrier appears between Possum Creek Spring (6), Brown Ranch (7) and Salamander Spring (9) on the east side of the barrier and western localities at Fessenden Springs (2), Hill Country SNA (3) and Western Kerr (4). Another barrier separates three proximal localities in the center of our sampling (Guadalupe River State Park (23), Honey Creek (24) and Preserve Cave (25)) from more westerly localities (Possum Creek Spring (6), Brown Ranch (7) and Salamander Spring (9)). In general, the estimated effective migration surface suggests that there are clear deviations from a simple IDB pattern across the sample range of this study which supports the hypothesis that there has been some gene flow during divergence or since populations and lineages have diverged.

**Figure 10:**
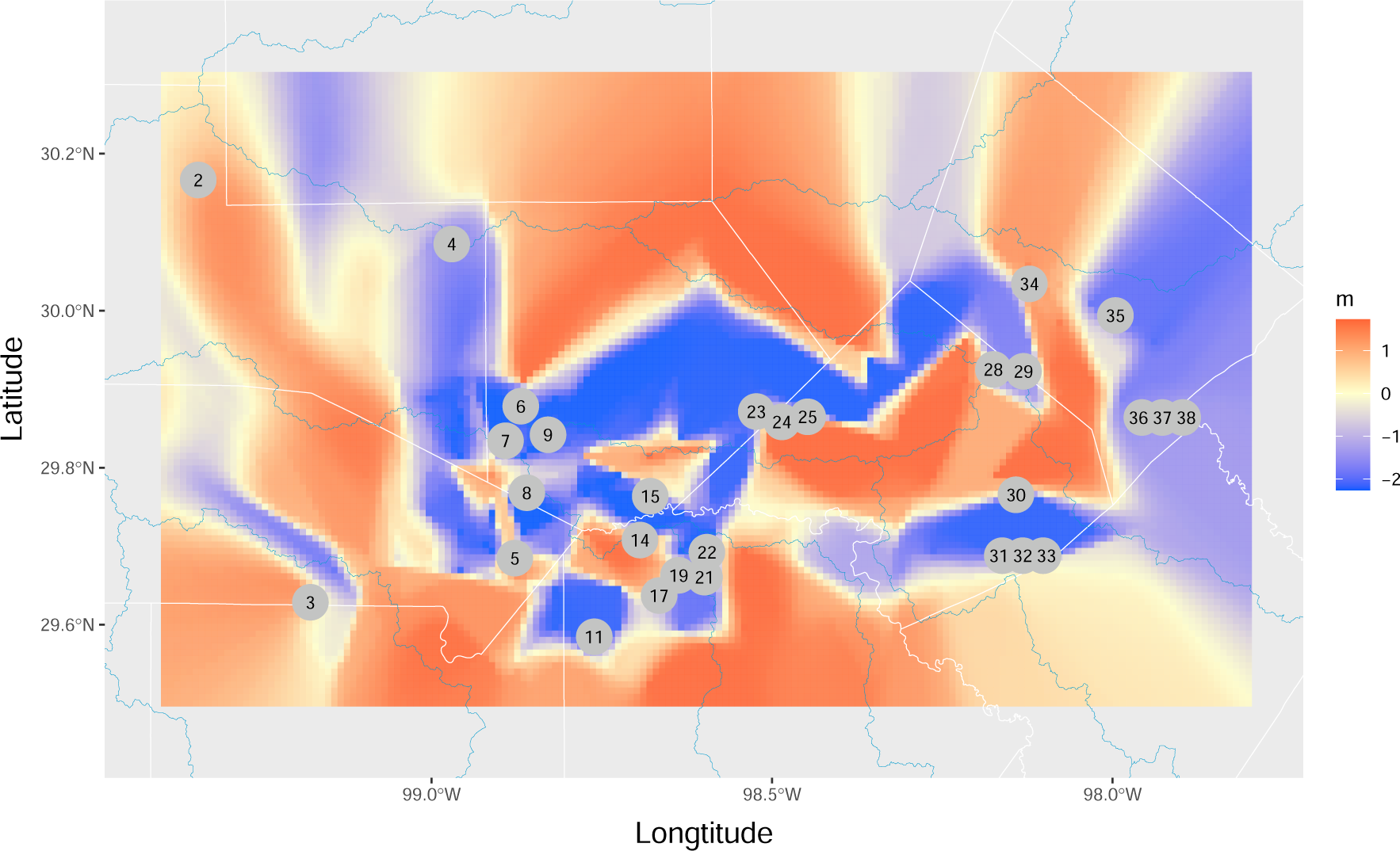
Estimated effective migration surface. The migration surface indicates departures from a simple Isolation-By-Distance, stepping stone model. Areas in red indicate higher migration (*m*) than expected; areas in blue indicate lower migration rates than expected. Locality numbers follow Table 1.

## Discussion

The history of salamanders of the *E. neotenes* complex (*E. neotenes*, *E. latitans* and *E. pterophila*) appears to be one of recent diversification among three lineages whose boundaries are obscured by substantial admixture not only among localities we sampled within the species complex, but also with adjacent lineages including *E. nana*, *E. troglodytes* and *E.* sp. 2 (sensu Devitt *et al*. (2019)) (Figures 3, 4). These patterns are consistent with a history of recent divergence accompanied by gene flow.

This conclusion is supported by several lines of evidence: First, differentiation among localities and taxa (measured as *F_ST_*) is low (Figures 2, 7, Supplementary Tables 1-4). Second, clustering analyses reveal prevalent admixture among and between localities and nominal species (Figure 3). Third, nominal taxonomy explains slightly more genetic variation than did geography in an RDA, but overall variance explained was low (Figure 6). Moreover, both taxonomy and geography were important predictors in Bayesian mixed models, but geography alone had greater support than taxonomy alone (Table 3). Fourth, analyses of admixture events revealed evidence of extensive gene exchange both within and among lineages (Figures 9, 4). These patterns of gene exchange suggest the *E. neotenes* salamanders are, or have been, capable of movement within at least some portions of the Edwards Plateau aquifers. However, while the addition of migration events in the treemix analyses improved our ability to explain variation in allele frequencies, that improvement was small (Figure 9) and it is important to note that the drift tree alone (*m* = 0) explained considerable variance. This indicates that some of the observed admixture might be derived from the recency of divergence and the retention of ancestral polymorphism. Fifth, negative *f*_3_ statistics for all sampling localities supports the inference of gene exchange. Sixth, examination of the estimated effective migration surface shows departures from a simple isolation-by-distance pattern which might be expected under a primary divergence scenario. Collectively, these lines of evidence support the hypothesis that the population genetic variation of the *E. neotenes* complex salamanders has been shaped by recent divergence, retention of ancestral polymorphism, and a history of gene exchange.

**Table 3:** Comparison of competing models partitioning population genomic variance. Shown are results of Bayesian linear mixed model regression ranked by Deviance Information Criteria (DIC). Coefficients are posterior means with 95% highest density intervals in parentheses. Models with “All sites” include the 29 sites with *n ≥* 8. Models with “Focal sites” include the 23 sites from the *E. neotenes* complex with *n ≥* 8.

| Model | Coefficient | Mean (95% HDI) | DIC |
| --- | --- | --- | --- |
| All sites - Taxonomy And Geography | intercept | -0.001 (-0.321, 0.338) | 485.1 |
| | $\beta_{TAX}$ | 0.386 (0.256, 0.521) | |
| | $\beta_{GEO}$ | 0.253 (0.201, 0.304) | |
| All sites - Full | intercept | 0.002 (-0.321, 0.326) | 487.0 |
| | $\beta_{TAX}$ | 0.388 (0.263, 0.520) | |
| | $\beta_{GEO}$ | 0.251 (0.193, 0.306) | |
| | $\beta_{AQU}$ | 0.012 (-0.101, 0.125) | |
| All sites - Geography | intercept | -0.004 (-0.384, 0.356) | 514.8 |
| | $\beta_{GEO}$ | 0.342 (0.297, 0.388) | |
| All sites - Geography And Aquifer | intercept | 0.023 (-0.321, 0.359) | 516.9 |
| | $\beta_{GEO}$ | 0.340 (0.290, 0.389) | |
| | $\beta_{AQU}$ | 0.008 (-0.115, 0.127) | |
| All sites - Taxonomy and Aquifer | intercept | -0.006 (-0.352, 0.335) | 561.9 |
| | $\beta_{TAX}$ | 0.704 (0.582, 0.827) | |
| | $\beta_{AQU}$ | 0.186 (0.077, 0.312) | |
| All sites - Taxonomy | intercept | 0.001 (-0.346, 0.335) | 568.3 |
| | $\beta_{TAX}$ | 0.749 (0.626, 0.865) | |
| All sites - Aquifer | intercept | 0.000 (-0.348, 0.362) | 676.3 |
| | $\beta_{AQU}$ | 0.344 (0.213, 0.474) | |
| Focal sites - Full | intercept | -0.002 (-0.208, 0.183) | 217.4 |
| | $\beta_{TAX}$ | 0.208 (0.091, 0.327) | |
| | $\beta_{GEO}$ | 0.237 (0.178, 0.301) | |
| | $\beta_{AQU}$ | -0.186 (-0.327, 0.046) | |
| Focal sites - Taxonomy and Geography | intercept | -0.001 (-0.177, 0.191) | 225.6 |
| | $\beta_{TAX}$ | 0.218 (0.104, 0.344) | |
| | $\beta_{GEO}$ | 0.204 (0.151, 0.264) | |
| Focal sites - Geography and Aquifer | intercept | -0.001 (-0.194, 0.201) | 229.7 |
| | $\beta_{GEO}$ | 0.295 (0.241, 0.348) | |
| | $\beta_{AQU}$ | -0.200 (-0.351, -0.056) | |
| Focal sites - Geography | intercept | 0.001 (-0.193, 0.184) | 238.7 |
| | $\beta_{GEO}$ | 0.263 (0.215, 0.308) | |
| Focal sites - Taxonomy | intercept | -0.008 (-0.198, 0.196) | 272.4 |
| | $\beta_{TAX}$ | 0.4641 (0.356, 0.579) | |
| Focal sites - Taxonomy and Aquifer | intercept | -0.003 (-0.192, 0.194) | 273.9 |
| | $\beta_{TAX}$ | 0.459 (0.348, 0.569) | |
| | $\beta_{AQU}$ | 0.034 (-0.103, 0.177) | |
| Focal sites - Aquifer | intercept | -0.002 (-0.179, 0.183) | 339.2 |
| | $\beta_{AQU}$ | 0.144 (-0.0014, 0.290) | |

Both the PCA and clustering analysis provide important insights into the organization of genetic variation within and among localities and lineages. While three loosely organized clusters are apparent within the *E. neotenes* complex (Figures 2, 3), genomic differentiation among localities and nominal taxa is low (Supplementary Tables 1-4) and many localities exhibit admixture (Figures 3, 4). Admixture is evident in localities such as Cascade Caverns (15), the type locality for *E. latitans* (Smith & Potter, 1946). Many localities exhibit admixture from multiple ancestral lineages, with Preserve Cave (25) showing admixture from five clusters (though the contribution from the *E. nana* cluster is minor). Specific tests of gene exchange using the treemix algorithm, and calculation of *f*_3_ statistics, indicate gene flow and patterns of allele frequency variation among localities that are not consistent with bifurcating relationships. Thus, a major finding of our research is that the three nominal taxa that have historically been recognized are not clearly differentiated. The lack of morphological characters available to differentiate among these species (Chippindale *et al*., 2000; Sweet, 1984), combined with evidence of extensive admixture among populations and lineages, indicate that the nominal species *E. neotenes*, *E. latitans*, and *E. pterophila* do not represent three independent lineages, but rather form three closely related management units of a single lineage with with a history of gene exchange among them.

Despite the evidence of admixture among *E. neotenes* localities, levels of genetic diversity (*θ* and *π*, Figure 5) were marginally lower across the complex compared to populations to the east (*E. nana*) and west (*E. troglodytes* and *E. sp. 2*) and two to three times lower than observed previously in *E. chisholmensis* (Nice *et al*., 2021). These low levels suggest that contemporary effective population sizes might be comparatively low in *E. neotenes* complex populations.

The inferred patterns of gene exchange provide clues to how *Eurycea* salamanders are, or have been, capable of moving through the aquifers such that the aquifers serve as a conduit for gene exchange. It is important to note that the analyses presented here cannot discriminate between gene flow that might have occurred during the divergence of the three admixed lineages that we observe in the *E. neotenes* complex and post-divergence gene flow. More detailed, explicitly demographic models are probably required for this, but might be difficult to apply given the scale of the sampling and the low levels of differentiation among sampled localities. Consequently, the details of this gene exchange and salamander movements remain obscure. What is evident by mapping the inferred migration events (Figure 4) is that gene flow was likely dynamic. Some of the migration events inferred from treemix analysis occurred along the path of known movement of water in the aquifer system in the central Edwards Plateau, which is thought to be generally and primarily from northwest to southeast within and across the constituent aquifers (Maclay, 1995). For example, inferred migration from Fesseneden Springs (2) to Western Kerr (4) and from Western Kerr (4) to Cascade Cavern (15) parallels known aquifer flows. However, other migration events do not align with the direction of water flow. For example, inferred migrations from *E. nana* (36, 37, 38) to the *E. neotenes* complex, particularly the southwestern localities (11, 17, 19, 21, 22), is nearly in the opposite direction, as is inferred migration from southwestern *E. neotenes* (11, 17, 19) to *E. troglodytes* (3), and from southwestern *E. neotenes* (17, 19) to Osborn Springs (5). These events suggest that salamander movements via the aquifers might have been dynamic over their history. Major fluctuations in aquifer water levels or unrecognized flow path changes might explain some of these patterns. Alternatively, the patterns we have elucidated might suggest that *Eurycea* movements are not constrained by aquifer water flow paths. However, it is also important to note that pathways for salamanders in the aquifers are undoubtedly quite complex and arrows plotted on a surface map (Figure 4) most assuredly represent an incomplete picture.

Similarly, inferred migration events from treemix also provide an incomplete picture of the history of gene exchange in these salamanders. Eight migration events inferred from treemix analysis explained a small proportion of the variance in allele frequencies among populations (Figure 8) but likely does not represent the full picture of gene flow. There were likely many more events, perhaps of smaller magnitude in terms of numbers of migrants or distance of migrations, that were not detected. The migration events inferred from treemix do not fully comport with the more extensive patterns of admixture observed in clustering analysis (Figure 3), which suggests that even more migrations have occurred in the history of these populations or that some of the admixture occurred during divergence. In addition, replicate runs for each model produced a range of different solutions (Figure 8) perhaps suggesting that many small migration events are also compatible with the data. Indeed, *f*_3_ statistics indicate that gene flow was detectable in all localities, emphasizing again that strictly bifurcating relationships are insufficient to capture the history of divergence at all localities.

The results reported here magnify previous work. Chippindale *et al*. (2000) used 22 allozyme loci and one mitochndrial DNA (mtDNA) locus to survey genetic variation across central Texas *Eurycea* and documented especially low-levels of differentiation among the *E. neotenes* complex taxa and localities, similar to results reported here. However, Chippindale *et al*. (2000) deemed the observed differentiation to be of sufficient magnitude to support delineating the nominal species within the *E. neotenes* complex (*E. neotenes*, *E. pterophila*, *E. latitans*, and *E. tridentifera*). Using mtDNA and a single-copy nuclear marker, Lucas *et al*. (2009) found modest amounts of sequence divergence among six localities from the eastern portion of the *E. neotenes* complex, though levels of differentiation estimated as *ϕ_ST_* (Excoffier *et al*., 1992) indicated more substantial allele frequency variation than reported here (pairwise *ϕ_ST_* ranged from 0.249 to 0.924 (Lucas *et al*., 2009)). These differences might reflect the limitations of a small number of sites or loci and the types of markers used by the previous studies.

Bendik *et al*. (2013) used mtDNA for a phylogenetic exploration of *Eurycea* south of the Colorado River, including the *E. neotenes* complex. They concluded that while there was clear structure, there was also evidence of admixed populations (especially compared to morphology) resulting from incomplete lineage sorting or gene exchange.

To date, the most comprehensive survey of genomic variation in central Texas *Eurycea* was presented by Devitt et al. (2019). These authors combined phylogenetic analyses of all central Texas *Eurycea* and clustering analyses of the southeastern group, including the *E. neotenes* complex, and concluded that the taxonomic boundaries among the *E. neotenes* complex species were not clear and advocated for recognizing fewer species in this region (Devitt *et al*., 2019). The current study expanded sampling of the *E. neotenes* complex, both in terms of number of localities and numbers of samples per locality. This expanded sampling has facilitated a fine-scale examination of geographic patterns of genomic variation and enabled identification of admixed localities and quantification of patterns of gene exchange. The level of admixture among seemingly isolated populations of salamanders observed at springs and in caves is surprising but highlights the potential connectivity of populations of the larger community of stygobionts over relatively large areas and over time. Comparative investigations of gene exchange will be needed to determine whether and how other stygobionts utilize the aquifer connections and whether life history variation predicts levels of connectivity. Our results also suggest that conserving the underground connections between populations of stygobionts might be important for ensuring the persistence of these aquatic populations in the face of increasing human impacts.

## Supporting information

Nice_et_al._SI

## Acknowledgments

We thank Travis LaDuc at the Texas Natural History Collections, Texas Memorial Museum at The University of Texas at Austin, and Tom Devitt for providing critical samples. We especially thank numerous land owners for access to springs and cave sites. We appreciate the cooperation from the US Air Force for providing access to historic localites. We thank Tom Devitt and David Hillis for constructive discussion. Sequencing was performed by the Genomic Sequencing and Analysis Facility at UT Austin, Center for Biomedical Research Support (RRID:SCR 021713). Computing was performed at the LEAP High Performance Computing Cluster at Texas State University. We thank Texas Parks and Wildlife for funding support through a Conservation License Plate Grant and the Texas Conservation Field Office for support with funding lab supplies. The views expressed in this paper are the authors’ and do not necessarily reflect the views of the U.S. Fish and Wildlife Service or Texas Parks and Wildlife Department.

## Author Contributions

CCN, PC, PHD conceived and designed the study. All authors collected the data. CCN, KLB and ZG analyzed the data. CCN drafted the initial version of the manuscript. All authors contributed to later versions of the manuscript.

## Data Availability Statement

The DNA sequence data analysed in this manuscript have been archived on NCBI’s SRA (PRJNA1057889). Genotype estimates and a predictors file for Redundancy Analysis and BLMM have been archived on Dryad (https://).

