## Supplementary material for "Extensive admixture among karst-obligate salamanders reveals evidence of recent divergence and gene exchange through aquifers": Nice_et_al._SI

Running title: *Population Genetics of* Eurycea *salamanders*

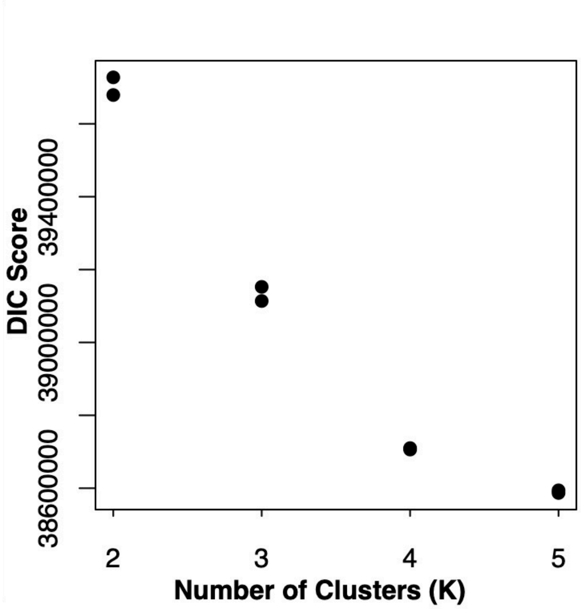


Supplementary Figure 1: Deviance information criterion scores for MCMC chains for ENTROPY models for k = 2 to 5.

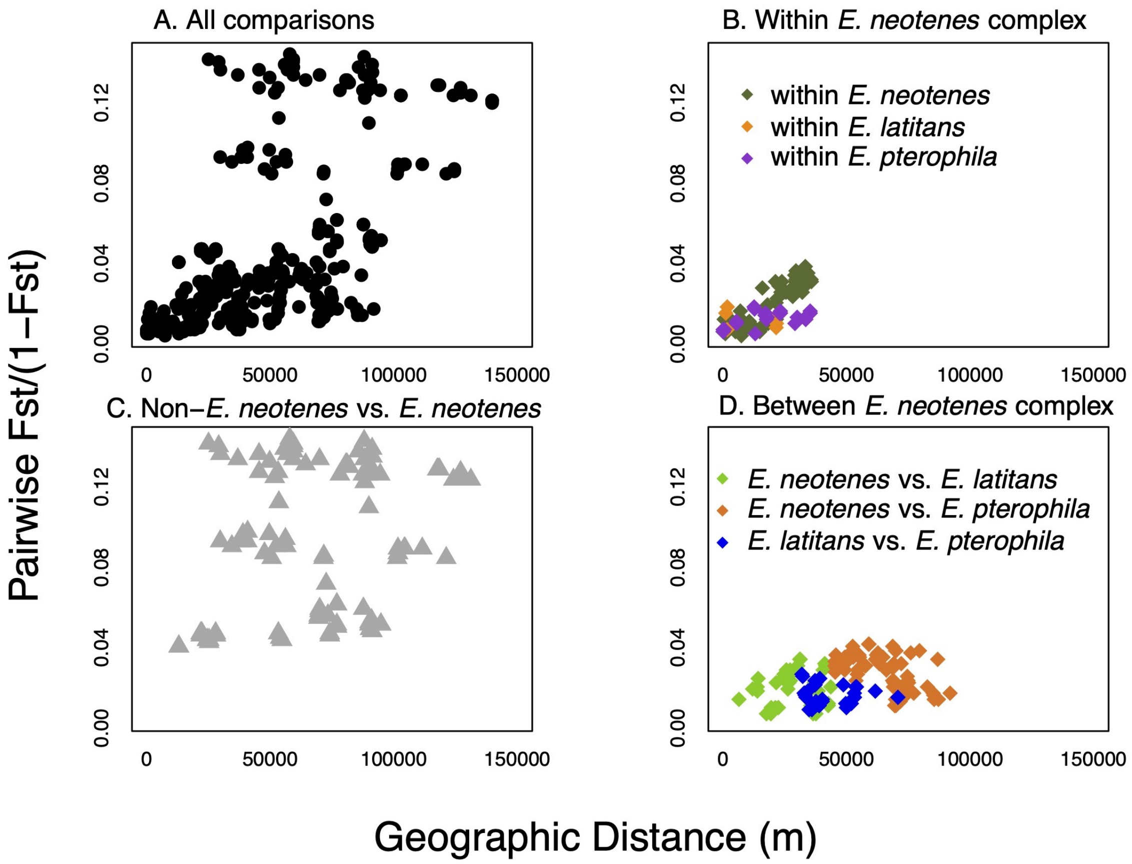


Supplementary Figure 2: Patterns of genetic differentiation in relation to geographic distances. A) Patterns of isolation by distance across all localities with $n \geq 8$. B) Patterns within the three nominal species in the *E. neotenes* complex. C) Patterns in comparisons of *E. neotenes* complex taxa versus non-*E. neotenes* lineages. D) Patterns in comparisons between *E. neotenes* complex taxa.

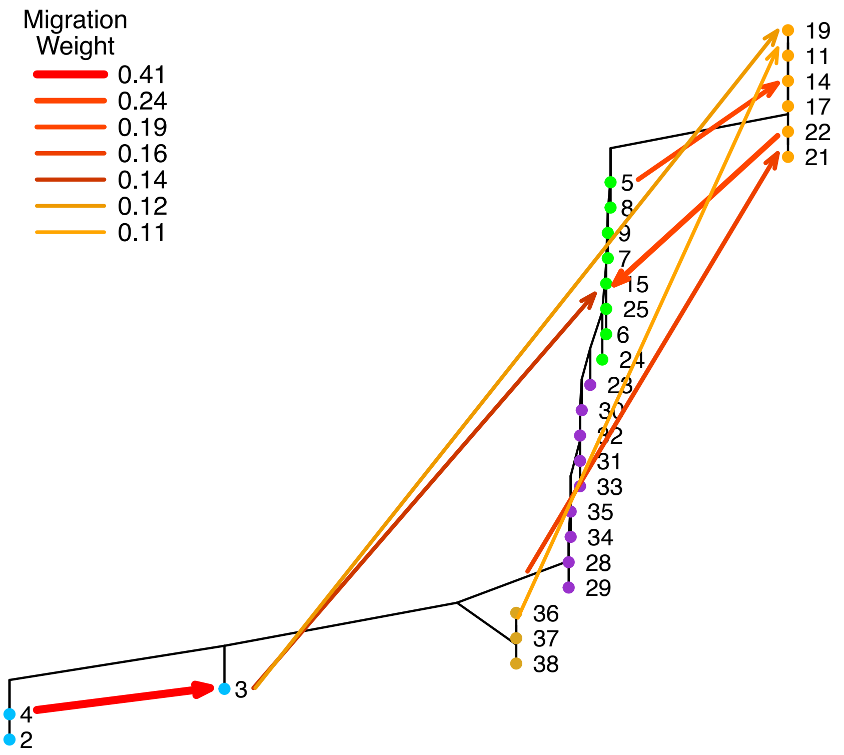


Supplementary Figure 3: Treemix tree of salamander localities with seven migration events (m = 7). Locality numbers follow Table 1. Locality 2 serves as the outgroup.

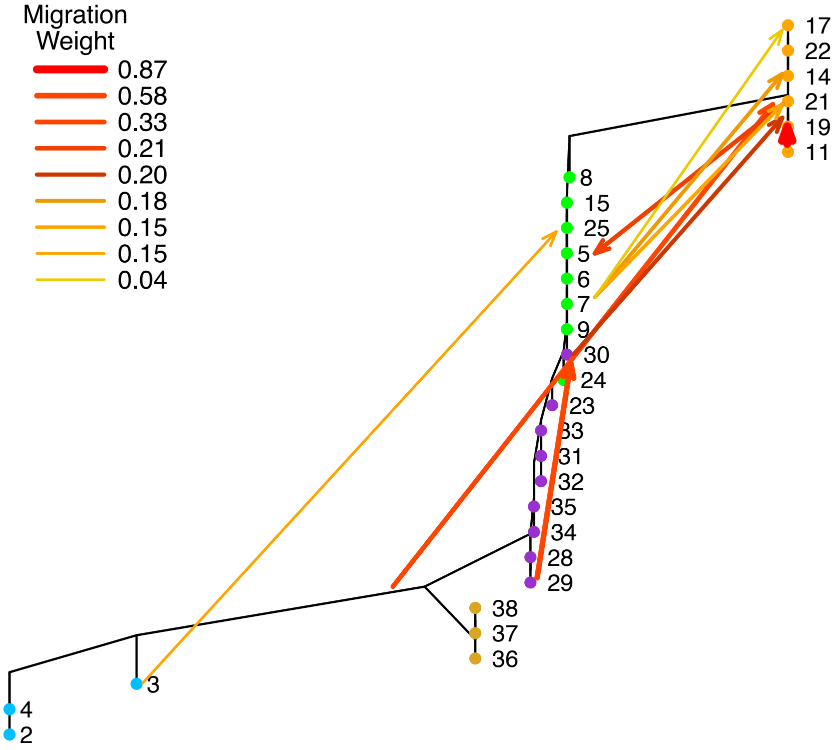


Supplementary Figure 4: Treemix tree of salamander localities with nine migration events (m = 9). Locality numbers follow Table 1. Locality 2 serves as the outgroup.

Supplementary Table 1: Pairwise $F_{ST}$ values among all sites with sample sizes of at least 8 individuals (below diagonal) and 95% permutational confidence intervals (above diagonal). Site numbering follows Table 1 and Figure 1

| **site:** | **2** | **3** | **4** | **5** | **6** | **7** | **8** | **9** |
|---|---|---|---|---|---|---|---|---|
| **2** | 0 | 0.031-0.034 | 0.010-0.011 | 0.114-0.123 | 0.117-0.126 | 0.114-0.122 | 0.112-0.121 | 0.116-0.126 |
| **3** | 0.032 | 0 | 0.033-0.036 | 0.081-0.088 | 0.085-0.092 | 0.081-0.088 | 0.080-0.087 | 0.085-0.091 |
| **4** | 0.011 | 0.034 | 0 | 0.116-0.125 | 0.119-0.128 | 0.115-0.124 | 0.113-0.122 | 0.118-0.127 |
| **5** | 0.118 | 0.085 | 0.12 | 0 | 0.011-0.012 | 0.009-0.010 | 0.008-0.008 | 0.012-0.013 |
| **6** | 0.122 | 0.089 | 0.124 | 0.012 | 0 | 0.006-0.007 | 0.007-0.008 | 0.009-0.009 |
| **7** | 0.118 | 0.085 | 0.12 | 0.009 | 0.006 | 0 | 0.004-0.004 | 0.005-0.006 |
| **8** | 0.116 | 0.083 | 0.118 | 0.008 | 0.008 | 0.004 | 0 | 0.006-0.007 |
| **9** | 0.121 | 0.088 | 0.123 | 0.012 | 0.009 | 0.005 | 0.007 | 0 |
| **11** | 0.121 | 0.085 | 0.123 | 0.027 | 0.034 | 0.031 | 0.028 | 0.034 |
| **14** | 0.115 | 0.08 | 0.117 | 0.019 | 0.025 | 0.02 | 0.018 | 0.022 |
| **15** | 0.112 | 0.078 | 0.113 | 0.01 | 0.01 | 0.007 | 0.007 | 0.01 |
| **17** | 0.125 | 0.088 | 0.126 | 0.03 | 0.037 | 0.033 | 0.03 | 0.035 |
| **19** | 0.118 | 0.083 | 0.12 | 0.025 | 0.032 | 0.028 | 0.025 | 0.03 |
| **21** | 0.122 | 0.086 | 0.124 | 0.026 | 0.031 | 0.027 | 0.025 | 0.029 |
| **22** | 0.119 | 0.083 | 0.121 | 0.024 | 0.03 | 0.025 | 0.023 | 0.027 |
| **23** | 0.109 | 0.078 | 0.111 | 0.012 | 0.011 | 0.009 | 0.009 | 0.011 |
| **24** | 0.112 | 0.079 | 0.113 | 0.011 | 0.009 | 0.007 | 0.007 | 0.009 |
| **25** | 0.099 | 0.067 | 0.101 | 0.02 | 0.022 | 0.019 | 0.018 | 0.022 |
| **28** | 0.114 | 0.082 | 0.116 | 0.023 | 0.022 | 0.02 | 0.02 | 0.023 |
| **29** | 0.114 | 0.082 | 0.116 | 0.022 | 0.022 | 0.02 | 0.019 | 0.023 |
| **30** | 0.11 | 0.078 | 0.112 | 0.014 | 0.013 | 0.011 | 0.011 | 0.014 |
| **31** | 0.113 | 0.082 | 0.115 | 0.018 | 0.017 | 0.015 | 0.015 | 0.018 |
| **32** | 0.111 | 0.08 | 0.113 | 0.017 | 0.016 | 0.013 | 0.013 | 0.016 |
| **33** | 0.112 | 0.08 | 0.113 | 0.016 | 0.016 | 0.013 | 0.013 | 0.016 |
| **34** | 0.114 | 0.082 | 0.115 | 0.02 | 0.019 | 0.017 | 0.017 | 0.02 |
| **35** | 0.11 | 0.078 | 0.112 | 0.017 | 0.016 | 0.014 | 0.014 | 0.017 |
| **36** | 0.107 | 0.079 | 0.11 | 0.049 | 0.05 | 0.047 | 0.046 | 0.049 |
| **37** | 0.108 | 0.08 | 0.11 | 0.049 | 0.051 | 0.047 | 0.046 | 0.05 |
| **38** | 0.108 | 0.08 | 0.11 | 0.049 | 0.05 | 0.047 | 0.046 | 0.049 |

Supplementary Table 2: Pairwise $F_{ST}$ values (continued) from Supplementary Table 1. Pairwise $F_{ST}$ values among all sites with sample sizes of at least 8 individuals (below diagonal) and 95% permutational confidence intervals (above diagonal). Site numbering follows Table 1 and Figure 1

| **site:** | **11** | **14** | **15** | **17** | **19** | **21** | **22** | **23** |
|---|---|---|---|---|---|---|---|---|
| **2** | 0.116-0.125 | 0.111-0.120 | 0.108-0.116 | 0.120-0.129 | 0.114-0.123 | 0.118-0.126 | 0.115-0.123 | 0.105-0.113 |
| **3** | 0.081-0.088 | 0.077-0.083 | 0.075-0.081 | 0.085-0.092 | 0.079-0.086 | 0.083-0.089 | 0.080-0.086 | 0.075-0.081 |
| **4** | 0.118-0.127 | 0.113-0.121 | 0.109-0.117 | 0.122-0.131 | 0.116-0.124 | 0.120-0.128 | 0.116-0.125 | 0.107-0.115 |
| **5** | 0.026-0.028 | 0.018-0.020 | 0.010-0.010 | 0.029-0.032 | 0.023-0.026 | 0.025-0.027 | 0.023-0.025 | 0.012-0.013 |
| **6** | 0.032-0.036 | 0.024-0.026 | 0.010-0.011 | 0.036-0.039 | 0.030-0.033 | 0.030-0.032 | 0.028-0.031 | 0.010-0.011 |
| **7** | 0.029-0.033 | 0.019-0.021 | 0.007-0.007 | 0.031-0.034 | 0.026-0.030 | 0.026-0.028 | 0.024-0.026 | 0.008-0.009 |
| **8** | 0.027-0.030 | 0.017-0.019 | 0.006-0.007 | 0.029-0.032 | 0.024-0.027 | 0.024-0.026 | 0.022-0.024 | 0.008-0.009 |
| **9** | 0.032-0.036 | 0.021-0.023 | 0.009-0.010 | 0.033-0.036 | 0.028-0.032 | 0.028-0.030 | 0.026-0.028 | 0.011-0.012 |
| **11** | 0 | 0.008-0.008 | 0.021-0.023 | 0.010-0.011 | 0.005-0.006 | 0.013-0.014 | 0.010-0.011 | 0.029-0.032 |
| **14** | 0.008 | 0 | 0.013-0.015 | 0.010-0.010 | 0.006-0.006 | 0.011-0.011 | 0.008-0.008 | 0.021-0.023 |
| **15** | 0.022 | 0.014 | 0 | 0.023-0.025 | 0.018-0.020 | 0.019-0.021 | 0.017-0.019 | 0.010-0.011 |
| **17** | 0.01 | 0.01 | 0.024 | 0 | 0.008-0.009 | 0.015-0.016 | 0.012-0.013 | 0.032-0.035 |
| **19** | 0.006 | 0.006 | 0.019 | 0.009 | 0 | 0.011-0.012 | 0.008-0.008 | 0.027-0.029 |
| **21** | 0.013 | 0.011 | 0.02 | 0.016 | 0.011 | 0 | 0.011-0.012 | 0.027-0.029 |
| **22** | 0.01 | 0.008 | 0.018 | 0.012 | 0.008 | 0.012 | 0 | 0.025-0.028 |
| **23** | 0.031 | 0.022 | 0.01 | 0.033 | 0.028 | 0.028 | 0.026 | 0 |
| **24** | 0.028 | 0.019 | 0.008 | 0.03 | 0.025 | 0.026 | 0.024 | 0.007 |
| **25** | 0.03 | 0.023 | 0.015 | 0.033 | 0.027 | 0.028 | 0.026 | 0.018 |
| **28** | 0.037 | 0.03 | 0.02 | 0.04 | 0.035 | 0.036 | 0.034 | 0.016 |
| **29** | 0.037 | 0.03 | 0.02 | 0.04 | 0.034 | 0.036 | 0.034 | 0.015 |
| **30** | 0.031 | 0.023 | 0.012 | 0.034 | 0.028 | 0.029 | 0.027 | 0.009 |
| **31** | 0.036 | 0.028 | 0.017 | 0.039 | 0.034 | 0.035 | 0.032 | 0.012 |
| **32** | 0.034 | 0.026 | 0.015 | 0.037 | 0.031 | 0.032 | 0.03 | 0.011 |
| **33** | 0.034 | 0.026 | 0.015 | 0.037 | 0.031 | 0.032 | 0.03 | 0.011 |
| **34** | 0.037 | 0.028 | 0.018 | 0.039 | 0.034 | 0.035 | 0.033 | 0.013 |
| **35** | 0.033 | 0.025 | 0.015 | 0.036 | 0.031 | 0.031 | 0.029 | 0.01 |
| **36** | 0.056 | 0.048 | 0.044 | 0.058 | 0.053 | 0.055 | 0.052 | 0.042 |
| **37** | 0.056 | 0.049 | 0.045 | 0.058 | 0.053 | 0.056 | 0.053 | 0.042 |
| **38** | 0.056 | 0.048 | 0.044 | 0.058 | 0.053 | 0.055 | 0.052 | 0.042 |

Supplementary Table 3: Pairwise $F_{ST}$ values (continued) from Supplementary Table 2. Pairwise $F_{ST}$ values among all sites with sample sizes of at least 8 individuals (below diagonal) and 95% permutational confidence intervals (above diagonal). Site numbering follows Table 1 and Figure 1

| **sites:** | **24** | **25** | **28** | **29** | **30** | **31** | **32** | **33** |
|---|---|---|---|---|---|---|---|---|
| **2** | 0.107-0.116 | 0.095-0.103 | 0.110-0.118 | 0.109-0.118 | 0.106-0.114 | 0.109-0.117 | 0.107-0.115 | 0.107-0.116 |
| **3** | 0.076-0.082 | 0.065-0.070 | 0.079-0.085 | 0.079-0.085 | 0.075-0.081 | 0.079-0.085 | 0.077-0.083 | 0.077-0.083 |
| **4** | 0.109-0.117 | 0.097-0.105 | 0.112-0.120 | 0.111-0.120 | 0.107-0.116 | 0.111-0.120 | 0.109-0.117 | 0.109-0.118 |
| **5** | 0.010-0.011 | 0.020-0.021 | 0.022-0.024 | 0.021-0.023 | 0.013-0.014 | 0.018-0.019 | 0.016-0.017 | 0.016-0.017 |
| **6** | 0.009-0.010 | 0.021-0.022 | 0.021-0.023 | 0.021-0.022 | 0.012-0.013 | 0.017-0.018 | 0.015-0.016 | 0.015-0.016 |
| **7** | 0.006-0.007 | 0.018-0.020 | 0.019-0.021 | 0.019-0.021 | 0.010-0.011 | 0.015-0.016 | 0.013-0.014 | 0.013-0.014 |
| **8** | 0.006-0.007 | 0.018-0.019 | 0.019-0.021 | 0.019-0.020 | 0.010-0.011 | 0.015-0.016 | 0.013-0.014 | 0.013-0.014 |
| **9** | 0.009-0.010 | 0.021-0.022 | 0.022-0.024 | 0.022-0.024 | 0.013-0.014 | 0.018-0.019 | 0.016-0.017 | 0.016-0.017 |
| **11** | 0.027-0.030 | 0.028-0.031 | 0.036-0.039 | 0.035-0.039 | 0.029-0.033 | 0.034-0.038 | 0.033-0.036 | 0.032-0.036 |
| **14** | 0.018-0.020 | 0.022-0.024 | 0.028-0.031 | 0.028-0.031 | 0.022-0.024 | 0.027-0.029 | 0.024-0.027 | 0.024-0.027 |
| **15** | 0.008-0.008 | 0.014-0.015 | 0.019-0.021 | 0.019-0.021 | 0.011-0.012 | 0.016-0.017 | 0.014-0.015 | 0.014-0.015 |
| **17** | 0.029-0.032 | 0.031-0.034 | 0.038-0.042 | 0.039-0.042 | 0.033-0.035 | 0.037-0.041 | 0.035-0.039 | 0.035-0.039 |
| **19** | 0.024-0.027 | 0.026-0.028 | 0.033-0.036 | 0.033-0.036 | 0.027-0.030 | 0.032-0.035 | 0.030-0.033 | 0.030-0.033 |
| **21** | 0.024-0.027 | 0.027-0.029 | 0.035-0.038 | 0.035-0.037 | 0.028-0.030 | 0.033-0.036 | 0.031-0.034 | 0.031-0.034 |
| **22** | 0.022-0.025 | 0.025-0.027 | 0.033-0.035 | 0.032-0.035 | 0.026-0.028 | 0.031-0.034 | 0.029-0.031 | 0.029-0.032 |
| **23** | 0.007-0.007 | 0.017-0.018 | 0.015-0.017 | 0.015-0.016 | 0.009-0.009 | 0.012-0.013 | 0.010-0.011 | 0.010-0.011 |
| **24** | 0 | 0.014-0.015 | 0.017-0.018 | 0.016-0.018 | 0.009-0.009 | 0.013-0.014 | 0.011-0.012 | 0.011-0.012 |
| **25** | 0.015 | 0 | 0.025-0.027 | 0.024-0.026 | 0.017-0.019 | 0.022-0.024 | 0.020-0.022 | 0.020-0.022 |
| **28** | 0.017 | 0.026 | 0 | 0.006-0.007 | 0.012-0.013 | 0.015-0.017 | 0.015-0.016 | 0.015-0.016 |
| **29** | 0.017 | 0.025 | 0.007 | 0 | 0.012-0.013 | 0.015-0.016 | 0.014-0.015 | 0.014-0.016 |
| **30** | 0.009 | 0.018 | 0.013 | 0.012 | 0 | 0.011-0.012 | 0.010-0.010 | 0.010-0.010 |
| **31** | 0.013 | 0.023 | 0.016 | 0.015 | 0.011 | 0 | 0.006-0.007 | 0.007-0.007 |
| **32** | 0.012 | 0.021 | 0.015 | 0.015 | 0.01 | 0.006 | 0 | 0.006-0.006 |
| **33** | 0.011 | 0.021 | 0.015 | 0.015 | 0.01 | 0.007 | 0.006 | 0 |
| **34** | 0.014 | 0.024 | 0.018 | 0.017 | 0.013 | 0.016 | 0.015 | 0.015 |
| **35** | 0.012 | 0.021 | 0.015 | 0.014 | 0.01 | 0.013 | 0.012 | 0.012 |
| **36** | 0.043 | 0.045 | 0.045 | 0.044 | 0.041 | 0.045 | 0.044 | 0.044 |
| **37** | 0.043 | 0.045 | 0.045 | 0.045 | 0.042 | 0.045 | 0.044 | 0.044 |
| **38** | 0.043 | 0.045 | 0.045 | 0.045 | 0.042 | 0.045 | 0.044 | 0.044 |

Supplementary Table 4: Pairwise $F_{ST}$ values (continued) from Supplementary Table 3. Pairwise $F_{ST}$ values among all sites with sample sizes of at least 8 individuals (below diagonal) and 95% permutational confidence intervals (above diagonal). Site numbering follows Table 1 and Figure 1

| **sites:** | **34** | **35** | **36** | **37** | **38** |
|---|---|---|---|---|---|
| | 34 | 35 | 36 | 37 | 38 |
| **2** | 0.109-0.118 | 0.105-0.114 | 0.103-0.111 | 0.104-0.112 | 0.103-0.112 |
| **3** | 0.079-0.085 | 0.075-0.081 | 0.076-0.082 | 0.077-0.083 | 0.077-0.083 |
| **4** | 0.111-0.120 | 0.108-0.116 | 0.106-0.114 | 0.106-0.115 | 0.106-0.115 |
| **5** | 0.019-0.021 | 0.016-0.018 | 0.047-0.051 | 0.047-0.051 | 0.047-0.051 |
| **6** | 0.018-0.020 | 0.016-0.017 | 0.048-0.053 | 0.049-0.053 | 0.048-0.052 |
| **7** | 0.016-0.018 | 0.013-0.015 | 0.045-0.049 | 0.045-0.049 | 0.045-0.049 |
| **8** | 0.016-0.018 | 0.013-0.014 | 0.044-0.048 | 0.045-0.049 | 0.044-0.048 |
| **9** | 0.019-0.021 | 0.016-0.017 | 0.047-0.052 | 0.047-0.052 | 0.047-0.051 |
| **11** | 0.035-0.038 | 0.032-0.035 | 0.053-0.058 | 0.053-0.058 | 0.053-0.058 |
| **14** | 0.027-0.030 | 0.024-0.026 | 0.046-0.050 | 0.047-0.051 | 0.046-0.050 |
| **15** | 0.017-0.019 | 0.014-0.016 | 0.043-0.046 | 0.043-0.047 | 0.042-0.046 |
| **17** | 0.038-0.041 | 0.034-0.037 | 0.055-0.060 | 0.055-0.060 | 0.055-0.060 |
| **19** | 0.032-0.035 | 0.029-0.032 | 0.051-0.055 | 0.051-0.055 | 0.050-0.055 |
| **21** | 0.033-0.036 | 0.030-0.033 | 0.053-0.057 | 0.053-0.058 | 0.053-0.057 |
| **22** | 0.031-0.034 | 0.028-0.030 | 0.050-0.055 | 0.050-0.055 | 0.050-0.055 |
| **23** | 0.012-0.013 | 0.010-0.011 | 0.040-0.044 | 0.041-0.044 | 0.040-0.044 |
| **24** | 0.014-0.015 | 0.011-0.012 | 0.041-0.045 | 0.041-0.045 | 0.041-0.045 |
| **25** | 0.023-0.025 | 0.020-0.022 | 0.043-0.047 | 0.044-0.047 | 0.043-0.047 |
| **28** | 0.017-0.019 | 0.015-0.016 | 0.043-0.047 | 0.044-0.047 | 0.043-0.047 |
| **29** | 0.016-0.018 | 0.014-0.015 | 0.043-0.046 | 0.043-0.047 | 0.043-0.047 |
| **30** | 0.012-0.013 | 0.010-0.011 | 0.040-0.043 | 0.040-0.044 | 0.040-0.043 |
| **31** | 0.015-0.017 | 0.013-0.014 | 0.043-0.047 | 0.043-0.047 | 0.043-0.047 |
| **32** | 0.014-0.016 | 0.012-0.013 | 0.042-0.045 | 0.042-0.046 | 0.042-0.046 |
| **33** | 0.014-0.016 | 0.012-0.013 | 0.042-0.046 | 0.042-0.046 | 0.042-0.046 |
| **34** | 0 | 0.005-0.005 | 0.040-0.044 | 0.040-0.044 | 0.041-0.044 |
| **35** | 0.005 | 0 | 0.037-0.040 | 0.037-0.041 | 0.037-0.041 |
| **36** | 0.042 | 0.039 | 0 | 0.005-0.006 | 0.006-0.006 |
| **37** | 0.042 | 0.039 | 0.005 | 0 | 0.006-0.006 |
| **38** | 0.042 | 0.039 | 0.006 | 0.006 | 0 |
